# You Are What You Sync: Neural Synchronization Predicts Religious Affiliation

**DOI:** 10.64898/2025.12.03.692126

**Authors:** Yohay Zvi, Nitai Kerem, Yaara Yeshurun

## Abstract

Existing research has found that shared neural responses to naturalistic narratives are associated with shared understanding, especially among members of the same social group. In this fMRI study, we tested whether in-group neural synchronization solely reflects explicit shared understandings, or relates to other group characteristics. To do so, we compared the neural synchronization patterns among two distinct social groups: Religious (*N* = 21) and Secular (*N* = 21). Participants were scanned while watching short video clips containing religious or neutral content and then answered questions regarding their reactions to- and interpretation of the videos. Behavioral results did not reveal group differences in responses’ homogeneity: neither in the engagement and agreement while watching the videos, nor in the emotions and verbal reactions elicited by it. However, neuroimaging results revealed that religious participants exhibited strikingly higher in-group synchronization than secular, both for religious and neutral narratives. Remarkably, it was possible to predict individuals’ religious affiliation solely based on their in-group neural synchrony, with accuracy scores of up to 92%. This pattern was consistent across all neural networks, most prominently in the Default Mode Network (DMN), Control and Attention networks. It also emerged in the Salience network and Somatomotor regions, hinting at neural processes that may foster group cohesion and identification. We propose that increased neural synchrony in the Religious group was driven mainly by greater homogenous social structure, suggesting a novel perspective for interpreting the role of neural synchrony in group dynamics.

## Introduction

Neural synchronization is a marker for shared understanding between individuals that view the world in a similar manner. In this study we examine whether in-group neural synchronization is also a characterizing feature of a social group, that arises not only from shared understanding of its members, but also from its underlying structure. We explore this question within the context of religious affiliation.

When introducing the Intersubject Correlation (ISC) analysis, Hasson et al. (2004) demonstrated how neural activity tends to align across individuals in regions that process a shared stimulus. Further studies exemplified how neural synchrony in low-level regions marks shared processing of stimuli’ physical properties, whereas neural synchrony in high-level regions marks shared understanding of stimuli’ meaning (Ames et al., 2015; Honey et al., 2012; Lerner et al., 2011). Neural synchronization reveals not only *if* individuals understand content, but also *how* they understand it, i.e., their interpretation. When individuals either watched an abstract animation or listened to a verbal description of the same story, those who interpreted the narrative similarly had more similar neural responses, regardless of whether they watched or listened (Nguyen et al., 2019). Likewise, sharing a psychological perspective synchronizes brain activity during natural viewing (Lahnakoski et al., 2014), and experimentally steering participants toward opposite narrative interpretations yields neural patterns that converge within, but diverge between, interpretations (Yeshurun et al., 2017).

Neural synchronization patterns also correspond to individuals’ social and cultural background. While watching a sport match, supporters of rivaling football teams shared neural response in low-order regions, but in high-order regions emerged group-dependent neural synchronization (Andrews et al., 2019). Similar patterns of group-dependent neural synchrony were found in the cultural domain, where individuals with different family background listened to an audiobook about their cultural background (Hakonen et al., 2022), and in the political domain, where individuals with a shared ideology shared neural responses when watching polarizing political video clips, distinct from those holding opposing attitudes (de Bruin et al., 2023; Katabi et al., 2023; Leong et al., 2020; van Baar et al., 2021).

Taken together, previous literature suggests that collective understanding within a social group is manifested in a synchronized neural pattern, different from the other groups’ pattern. However, whether the magnitude of neural synchronization – and not the specific neural pattern – differs between social groups, remains an unexplored aspect. An instance for such difference was found in a recent study between right- and left-wing individuals while watching polarizing political video-clips. Right-wing participants were characterized with higher levels of neural synchronization in response to most of the videos (while an opposite pattern was observed for one of the four political videos), with partisanship-dependent differences in synchronization already in early sensory, somatosensory and motor regions (Katabi et al., 2023). These findings imply that different extents of neural synchronization can serve as a marker for social polarization, emphasizing its relevance in understanding social mechanisms.

Religion is a salient social–cultural marker that differentiates religious and non-religious people across cognitive and social domains. Religious individuals commonly adopt “meaning-making” perspectives (Inzlicht et al., 2011; Vishkin et al., 2016) and use religious explanations for everyday events (Lupfer et al., 1992). Compared with more open-minded, socially heterogeneous non-religious, religious exhibit lower cognitive flexibility (Uzarevic & Coleman, 2021; Zmigrod et al., 2019) and a more homogeneous social structure shaped by shared norms (Ruiter & van Tubergen, 2009), which is mirrored in uniform linguistic patterns (Yaden et al., 2018) and intergroup attitudes (Johnson et al., 2012).

In this functional Magnetic Resonance Imaging (fMRI) study, we explore whether religious and secular groups exhibit distinct patterns of neural synchronization when processing religiously sensitive content. We hypothesized that members within each group would show similar neural responses to such content — reflecting a shared in-group interpretation — but that these neural patterns would differ between the religious and secular groups, with the religious, given the more homogenous social structure, exhibiting higher level of similarity in their neural responses.

## Methods

### Participants

Forty-two participants (27 female and 15 male) between the ages of 21 to 33 (*M* = 25.0, *SD* = 3.16) participated in the study. All participants were healthy, right-handed, had normal or corrected to normal vision, and met the health criteria required to participate in an MRI study (e.g., no tattoos above the neck). All participants were fluent in Hebrew at a mother tongue level. They were recruited through social networks as well as through “snowballing sampling” where participants recruited others upon completing the experiment themselves. All participants were screened through a preliminary online questionnaire (see *Supplementary Materials*) and allocated to the appropriate group according to their responses. We recruited only respondents who reported belonging to one of the following groups (1) religious (13 female and 8 male, age *M* = 25.2, *SD* = 3.14) or (2) Secular (14 female and 7 male, age *M* = 24.8, *SD* = 2.19).

In the religious group, we only recruited participants who identified themselves as part of the Religious-Zionism sector. We chose to focus on participants from the Religious-Zionism sector because this group tends to engage more actively with secular life in Israel, making them more likely to be familiar with the stimuli presented in the experiment. Additionally, this choice helped prevent potential discomfort for participants from more orthodox sectors, who might find the stimuli inappropriate or offensive.

Each group contained 21 participants, a sample size that has been shown to be sufficient in previous studies testing for similarities and differences in neural responses to naturalistic stimuli (Yeshurun et al., 2017; Katabi et al, 2023), and in a power analysis for the fMRI analysis method we used (Pajula & Tohka, 2016).

The study was approved by Tel Aviv University’s ethics committee and the Helsinki Institutional Committee of the Sheba Tel-Hashomer Medical Center. All participants provided written informed consent to participate in the study and received monetary compensation for their time.

### Stimuli and Experimental Design

Participants watched 3 video clips inside the MRI scanner: (i) a news report about the weather (2:07 minutes), which is neutral in terms of religious narrative (Neutral); (ii) a manifest of a religious person explaining about the dangers of alternative Kosher organizations (as opposed to the institutionalized Rabbinic organization which is generally more conservative) (2:23 minutes) (Kosher); and (iii) a news report of hospitals in Israel banning visitors’ Chametz (foods which are religiously forbidden to eat during Passover according to the Jewish tradition) on account of the hospitals being observant and Kosher (2:33 minutes) (Passover).

Each clip was preceded and followed by a grey screen: 8 seconds before, and 10 seconds after (which were discarded from the analysis). After each video-clip, participants were asked to answer three questions about it: (1) “How much did you agree with the main message of this clip?”; (2) “How much did this clip interest you?”; and (3) “How emotionally engaged were you?”. Participants answered these questions by indicating their ratings (using a magnet-compatible mouse) on a Visual Analog Scale (VAS) that ranged from “Not at all” (0) to “Very” (100). The first video was always the Neutral video, while the other two videos appeared in a predetermined random order.

Following the video task, participants also performed a recognition task which we did not include as part of this study.

During the entire fMRI scan, participants’ eye gaze was monitored and recorded at 500 Hz using SR-Research EyeLink 1000 Plus eye-tracker. However, due to technical issues, eye-tracking data was not collected for most participants, thus we did not analyze it.

Immediately following the scan, participants completed a behavioral assessment session in a separate room within the vicinity. In this session, they were asked to rate their emotional response towards each video regarding six explicit emotions (happiness, sadness, fear, disgust, anger and surprise) on a 1-5 Likert scale. Additionally, participants were asked to describe in a free text what they have seen in the video and what was the first thing that popped up to their head after watching the video. The final part of the session was a demographic survey (see *Supplementary Materials)*.

### fMRI Data Acquisition

Participants were scanned using a 3T Siemens Prisma scanner with a 64-channel head coil at the Tel-Aviv University Strauss Center for Computational Neuroimaging. T1-weighted structural images were acquired using a magnetization prepared rapid gradient echo pulse sequence (MPRAGE) as following: TR = 2530ms, TE = 2.88ms, TI 33 = 1100ms, flip angle = 7° and 250Hz/px, isotropic voxel size of 1mm³. For Functional scans, images were acquired using a T2*-weighted multiband echo planar imaging protocol. Repetition time (TR) = 1000 ms, echo time (TE) = 34ms, flip angle (FA) = 60°, multiband acceleration factor of six without parallel imaging. Isotropic resolution was 2mm³ (no gaps) with full brain coverage; slice-acquisition order was interleaved.

## Imaging Analysis

### Preprocessing

Raw DICOM format imaging data was converted to NIfTI with the dcm2nii tool. The NIfTI files were organized according to the Brain Imaging Data Structure (BIDS) format v1.0.1. fMRI data preprocessing was conducted using the FMRIB’s Software Library’s (FSL v6.0.2) fMRI Expert Analysis Tool (FEAT v6.00) (Sm et al., 2004). All data was subjected to the following preprocessing procedures: brain-extraction for skull-stripping anatomy image; slice-time correction; high-pass filtering (two cycles per stimulus’ length); motion-correction to the middle time-point of each run; and smoothing with a 4-mm FWHM kernel. All images were registered to the high-resolution anatomical data using boundary-based reconstruction, and normalized to the Montreal Neurological Institute (MNI) template (Mazziotta et al., 2001), using nonlinear registration. Blood-oxygen level-dependent (BOLD) response was normalized (z-scored) within subjects for every voxel for each video-clip. Hemodynamic response function (HRF) was calculated for each participant according to the peak start time of the BOLD response in early auditory areas (A1+), using the Neutral video clip. The shift was then calculated as the duration from the stimulus onset to the first peak of the hemodynamic response in A1+ for each participant (Religious mean shift = 3.62 s, SD = 0.74 s and Secular mean shift = 3.38 s, SD = 0.8 s). We further analyzed the data only in voxels that had a reliable BOLD signal (< 3000 AU) in at least 80% of the participants in each group. This procedure resulted in 224,821 voxels for the Neutral video clip, 224,770 voxels for the Kosher video clip and 224,577 voxels for the Passover video clip.

## Data Analysis

### Behavioral Data: Scanner Ratings for Agreement and Engagement

We analyzed participants’ Visual Analog Scale (VAS) ratings (agreement, interest, emotional engagement) using nine two-tailed Mann–Whitney U tests, with group (Religious vs. Secular) as the independent variable. Mann–Whitney was chosen due to the relatively small sample size (*N* ≤ 21 in both groups; due to technical problems with the mouse, the VAS scores for two participants from the Religious group were not recorded) and potential non-normality. In addition, to examine whether one group exhibited greater in-group homogeneity in their ratings, we performed Levene’s test for equal variances on each measure, for each of the questions separately.

### Behavioral Data: Post fMRI Scan questionnaire

#### Explicit Emotional Response

For the post-scan questionnaire, participants rated the extent to which each of six explicit emotions were elicited by the videos. We conducted a separate Mann–Whitney U test for each emotion with group as the independent variable. Once again, we measured potential in-group homogeneity using Levene’s test for equal of variances, for each of the questions separately.

#### Text Analysis

#### LIWC

For the two free text questions in the post scan questionnaire, we performed a descriptive linguistic analysis. We used LIWC2015 software (Pennebaker et al., 2015), which extracts 89 psycholinguistic and syntactic features per entry, ranked on a 0-100 scale. Responses were categorized according to experimental condition, defined by a 3 (vide-clips: *Neutral, Kosher, Passover*) × 2 (question type: *association, description*) × 2 (group: *religious*, *secular*) design. LIWC outputs were compared between groups separately for each task × question condition. For each comparison, we computed a two-tailed independent-samples *t*-tests to assess differences in mean LIWC scores, and Levene’s tests to evaluate homogeneity of variance. For descriptive purposes, in each comparison separately we excluded features where 20% or more of the responses had zero values, except for core summary metrics (Analytic, Clout, Authentic, Tone), which were retained regardless of sparsity.

#### Cosine Similarity

We applied another approach to test the semantic alignment across individuals using a method adapted from Duran et al. (2019). In each of the six videos × question comparisons (Neutral, Kosher, Passover × Description vs. Association), participants’ free-text responses were loaded into a quanteda corpus, and tokenized after stripping punctuation, symbols, and numbers. We then built three document-feature matrices (DFMs) — one on raw tokens, one on lemmas (via a custom token → lemma lookup), and one on part-of-speech tags (via a token → POS lookup) — each trimmed to terms present in at least two documents and re-weighted by proportional term frequency × inverse document frequency (TF-IDF). For each DFM, we calculated the group’s mean feature vector and used quanteda.textstats to compute each response’s cosine similarity to that mean. We derived a fourth “semantic” metric by averaging pre-trained Twitter word embeddings (adopted from the Hebrew word2Vec GitHub project, https://github.com/Ronshm/hebrew-word2vec?tab?readme-ov-file) per document and measuring cosine similarity to the global mean embedding. All four sets of cosine scores were Fisher-z-transformed and subjected to independent-samples t-tests contrasting religious and secular respondents.

### In-group Intersubject Correlation

To measure the level of in-group synchrony level within the Religious and Secular groups, we used Intersubject Correlation (ISC) (Hasson et al., 2004). ISC measures the degree to which neural responses to naturalistic stimuli are shared between participants processing the same stimuli. In each voxel, we correlated each participant’s time course with the average time-course response across all other participants in the same group using Pearson correlation, in a Leave-One-Out (LOO) procedure. We then averaged these 21 correlation coefficients values (since *N* = 21 in both the Religious and Secular) to get an estimation of the in-group similarity in the neural response in each of the voxels. This procedure was performed separately in each group, for each video.

#### Determining a Statistical Significance Threshold of the ISC scores

To define in which voxels the synchrony level was above chance, we compared the in-group ISC scores of each voxel to a null distribution that was generated in a permutation procedure. We repeated the same LOO procedure as described before, only this time we shuffled the group’ empirical time series using a phase-randomization procedure. Phase randomization was performed by Fast Fourier Transform (FFT) of the signal, randomizing the phase of each Fourier component, and then inverting the Fourier transformation back to the time domain. This procedure leaves the power spectrum of the signal unchanged but removes temporal alignments of the signals. Then, Pearson correlation was computed between the average time series of the group’s shuffled time-series, and a participant’s intact empirical time-series.

We repeated this procedure 1,000 times, leaving us with six null distributions (3 video-clips × 2 groups) of Voxels × 1,000 iterations (null distributions’ statistical parameters are described in table S1a in *Supplementary Materials*). Since a null distribution of 1,000 iterations does not allow the *p*-value to get lower than 0.001, we fitted for each voxel a Cumulative Distribution Function (CDF) based on the mean and Standard Deviation (SD) parameters of the null distribution in that specific voxel. We then computed the *p*-values for the actual ISC score based on the CDF, in each voxel separately. Lastly, we applied Benjamini and Hochberg (BH) False Discovery Rate (FDR) correction for multiple comparisons (Benjamini & Hochberg, 1995), using *q* criterion of 0.0005. This left us with voxels that the ISC score in them was above chance, in each group and video separately.

For further analyses, we defined Voxels of Interest (VOIs) as voxels that had an above chance ISC score in at least one of groups, Religious or the Secular. The number of VOIs was 65,792 in the Neutral video, 94,254 in the Kosher video and 88,143 in the Passover video.

### In-group ISC comparison: Religious vs Secular

#### Unique vs Overlap Synchronized Brain Regions

To examine the differences in the in-group synchrony between the Religious and the Secular groups, we compared their ISC scores. First, we counted the number of voxels with above-chance ISC score in each group. Following that, we compared the voxels in both groups to distinguish between voxels in which the ISC was above chance only in the Religious (“unique Religious”), only in the Secular (“unique Secular”) or in both groups (“overlap”). This procedure was performed in each video separately.

In the unique voxels of both groups, we conducted a one-tail t-test between the ISC scores, hypothesizing higher synchronization in the group these voxels were unique. The t-test was followed by an FDR correction to control multiple comparisons with a standard *q* criterion of 0.05. To test for differences in the in-group synchrony in the overlapping regions, we conducted a two-tailed t-test in these voxels.

A Chi-squared two-proportions test was carried out (Laurie, 2025) for each of the comparisons described above, to test for differences between the voxels counts in both groups, followed by an FDR correction for multiple comparisons.

#### Support Vector Machine (SVM) Classification

We trained an SVM classifier (Cortes & Vapnik, 1995) to test whether participants’ religious affiliation (Religious or Secular) can be predicted based on the level of neural synchronization they share with their group. We did so for each of the three video clips (*N* = 21 in each group), in each VOI. This classifier was carried using a leave-two-out algorithm that received a training set and a testing set. We used one-dimensional space data, the ISC score of each participant to classify the participants’ religious affiliation (i.e., the correlation coefficient between each participant’s brain response to the video and the averaged brain response of the rest of the group). The training set contained *N*-1 correlation coefficients of each group, and the testing set contained the remaining two correlation coefficients (one from each group). The support vector classifier used the training dataset to find a single point to serve as the hyperplane that classified the data into classes, and labeled the testing set as religious or secular accordingly. This algorithm was executed 𝑁^2^ times (each time, two different participants were left out). At each time, the classifier could be correct (1, i.e., classifying both test data points correctly), chance level (0.5, i.e., classifying one text data point correctly) or incorrect (0, i.e., classifying both test data points incorrectly). The classifier accuracy in a specific VOI was the average of the 𝑁^2^ trials (i.e., a number between 0 and 1).

##### Determining a statistical significance threshold of the SVM accuracy rates

To test whether the classifier accuracy value was significantly larger than would be expected by chance, we simulated a null distribution using a permutation procedure. The data (21 in-group correlations coefficients for each group) from a specific voxel was extracted, and the labels of the groups were shuffled randomly to create two new pseudo-groups. Due to the heavy computation cost of repeating the same leave-two-out procedure performed on the real data for 1,000 iterations, we did not divide the data into training and testing sets; instead, we intentionally overfitted an SVM model that was trained on the entire dataset, and measured its accuracy (0-1, as described above). This maximizes the model’s performance by fitting to all available data, making it harder for random permutations to achieve similar accuracy. This approach bounds the threshold to be stringent, thus compensating for not repeating the exact same procedure done on the real data. We repeated that procedure 1,000 times for each of the videos separately, resulting in three null distributions simulating the accuracy scores expected under the null hypothesis. We set the 95^th^ percentile of each of the null distributions, which was 0.6429 (consistent in all the videos), as a threshold for above-chance classification accuracy, excluding voxels with lower classification accuracy from further analysis (null distributions’ statistical parameters are described in table S1b in *Supplementary Materials*).

To correct for multiple comparisons, we applied Cluster Size Correction. For each of the 1,000 iterations of the null distribution, we generated a NIfTI brain image. Using FSL’s ‘cluster’ function, we identified clusters of voxels with accuracy scores above the threshold (the 95th percentile of the null distribution, as previously described). Clusters were defined based on six-neighbor connectivity. This procedure allowed us to estimate the size of clusters expected to occur under the null hypothesis, resulting in a distribution of these expected cluster sizes. To decide the minimum cluster size to threshold the real data, we considered two parameters: (a) the percentile of the clusters distribution and (b) whether to include very small clusters (<4 voxels) in the clusters’ distribution. We applied a two-parameter tuning grid, in which we simulated the exclusion of clusters of the size of 1, 2 or 3 from the null distribution, and defined a linear space of ten values between 95 and 99.9 to sample different percentiles. When observing the tuning grid results, we identified a notable ‘kink’. This approach is akin to methods used in clustering and optimization (e.g., the elbow method) where a notable ’kink’ or knee point in the plot indicates an inflection point that serves as an optimal cutoff. The observed kink suggested the 99.356 percentile as the threshold, without excluding small clusters from the null distribution when computing the percentile (see Fig. S1 in *Supplementary Materials*). This means that in the results we only included voxels that had an accuracy score higher than 95^th^ percentile of the null distribution, and were part of a cluster that was at least equal in size to the 99.356 percentile of the clusters’ size null distribution (36 clusters in the Neutral video, with minimum cluster size of 36 voxels; 47 clusters in the Kosher video, with minimum cluster size of 41 voxels; 47 clusters in the Passover video, with minimum cluster size of 43 voxels).

##### Spatial Expansion of Voxel Classifications and Cross-Video Consistency

To investigate the consistency of regions where SVM classification exceeded chance across all three videos, we applied a spatial expansion technique to the voxels identified in each video. This approach allowed us to assess the spatial consistency of neural synchronization patterns across different stimuli, focusing on regions that reliably distinguished religious from secular participants, even when individual voxel overlaps were limited. Using a spherical structuring element (Kriegeskorte et al., 2006) with a dilation radius of 1, we expanded each voxel where classification accuracy was above chance. In the Neutral video task, the expansion increased the number of voxels from 7,627 to 19,729; in the Kosher video task, the expansion increased the number of voxels from 5,653 to 15,716; in the Passover video task, the expansion increased the number of voxels from 9,185 to 23,409. We performed two sets of intersections: between the two religious videos (Kosher and Passover) and between all three videos.

### Brainnetome Parcellation and Networks Division

Although all the analyses were performed in a voxel-by-voxel manner, to facilitate the interpretation of the results and test their robustness across different regions, we also grouped single voxels into gyri and brain networks. We used the Human Brainnetome Atlas (BNA) (Fan et al., 2016), a connectivity-based atlas with 210 cortical subregions and 36 subcortical subregions. The atlas follows Yeo’s division into 7 and 17 brain networks division (Thomas Yeo et al., 2011), as well as providing parcellation into 24 gyri. For each of the 17 networks, we computed the proportion of synchronized network by counting in each network the number of voxels demonstrating in-group neural synchrony, divided by the number of total voxels mapped to this network. We repeated this procedure separately both for the voxels that exhibited higher than chance in-group synchrony, as well as for the voxels where group differences were found after performing t-test and applying FDR correction for multiple comparisons. To measure group differences in the proportion of synchronized networks, we conducted a series of 102 Chi-squared two-proportions test (17 networks × 3 video clips × 2 proportions’ measures) followed by Bonferroni correction for multiple corrections.

## Results

### Similarities and Differences in Engagement between Religious and Secular

A series of independent two-tailed Mann-Whitney U tests was conducted to examine group differences between the Secular and Religious for agreement, interest and emotional engagement with the video clips (based on their rating inside the scanner).

Significant group differences were found only for the Kosher video: the Religious group reported higher levels of agreement, interest and emotional engagement compared to the Secular group (*M* = 47.50, *SD* = 32.00; *M* = 17.60, *SD* = 25.55, *p* = 0.001; *M* = 72.82, *SD* = 15.80; *M* = 48.97, *SD* = 28.32, *p* = 0.016*; M* = 77.52, *SD* = 15.12; *M* = 52.78, *SD* = 27.13, *p* = 0.003, respectively) (Fig.1a). No significant group differences were found for the neutral or the Passover videos.

Additionally, Levene’s test for equality of variances was conducted to assess group differences in terms of in-group homogeneity. The assumption of equal variances was met for all measures (*p* > 0.05), besides for the level of interest in the Kosher video. In this case, the variance was significantly smaller in the Religious group (*SD* = 15.80) compared to the Secular group (*SD* = 28.32), *F*(1, 38) = 8.90, *p* = 0.005 (full report of the results is described in table S2 in *Supplementary Materials*). Thus, in 8 (out of 9) ratings conducted inside the scanner, there were no group differences in how variable the ratings were between the secular and the religious groups.

### Similarities and Differences in Emotional Responses between Religious and Secular

A series of independent two-tailed Mann–Whitney U tests was conducted to examine group differences between the Secular and Religious groups in the extent to which six explicit emotions were elicited by the video clips. Significant group differences were found only for anger in the Passover video: the Religious group reported significantly lower levels of anger (*Median* = 2.00, *IQR* = 1.00) compared to the Secular group (*Median* = 4.00, *IQR* = 1.00) (see Fig. 2b).

For happiness, the Religious group ranked slightly higher than the Secular group in both the Kosher and Passover videos. In the Kosher video, the Religious group had a slightly higher mean happiness score (*Mean* = 1.29, *SD* = 0.56) compared to the Secular group (Mean = 1.00, *SD* = 0). Similarly, in the Passover video, the Religious group reported slightly higher levels of happiness (*Mean* = 1.43, *SD* = 0.81) compared to the Secular group (*Mean* = 1.00, *SD* = 0). Since all Secular participants rated happiness as 1 (with no variability), we did not perform a Mann–Whitney U test for these comparisons.

Additionally, Levene’s test for equality of variances was conducted to assess group differences in terms of in-group homogeneity. Due to the lack of variability in the Secular group’s happiness ratings in both the Passover and Kosher videos, group differences could not be assessed in these cases. For all other measures, the assumption of equal variances was met (*<u>p</u>* > 0.05), except for anger in the Neutral video, where the Religious group showed significantly smaller variance (*SD* = 0.51) compared to the Secular group (*SD* = 1.12), *F*(1, 40) = 11.34, *p* = .002 (full report of the results is described in table S3 in *Supplementary Materials*). Thus, in 15 (out of 18) ratings on the emotions elicited by the videos, there were no group differences in how variable the ratings were between the secular and the religious groups, in 2 ratings the variability within the secular group was 0, and in one additional rating the variability within the religious group was smaller than within the secular group.

### Text Analysis

#### >LIWC

We performed 89 (LIWC output features) × 2 (question type: association vs description) × 3 (video clips) t-test for group differences in the text responses. Out of these 534 comparisons, we identified a total of 17 cases with significant group differences (p < .05). No consistency was found, i.e., none of the features showed significant group difference more than once. Levene’s tests revealed significant variance differences in a total of 64 features and 113 cases, with variance being greater in the religious group in 71 cases, and in the secular group in 42 (Fig 1c) (full description is in figure S2 in *Supplementary Materials*).

**Fig. 1.**
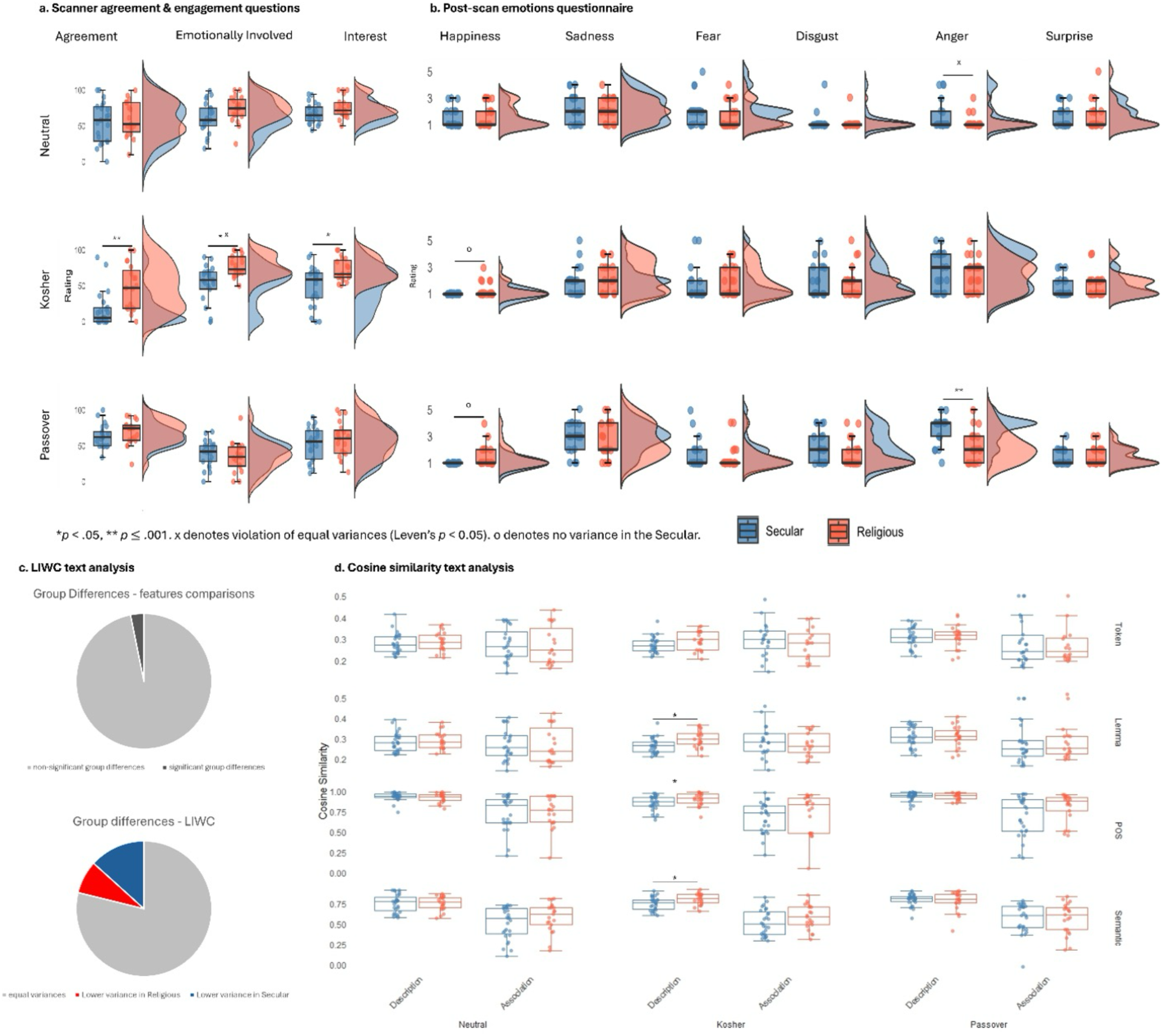
Behavioral ratings. **a**. **Visual Analog Scale (VAS) Results.** Participants were presented with three questions after each video presented, in which they answered during the fMRI scan on a scale of 0-100. Significant differences between the Religious and the Secular groups were found only towards the Kosher video, with the Religious reporting higher levels of agreement, interest and being more emotionally involved. Note: \**p* < .05, ** *p* ≤ .001; x denotes violation of equal variances (Leven’s *p* < 0.05**). b. Explicit Emotions Post-Scan questionnaire results.** Post scan results demonstrated high level of similarity in the explicit emotional responses between the Secular and Religious. Significant difference was found for the anger level in the Passover video. Due to the lack of variance in the Secular participants’ responses for the happiness levels in the Passover and Kosher video, we did not perform statistical test in these cases. Note: ** *p* ≤ .001; x denotes violation of equal variances (Leven’s *p* < 0.05); o denotes no variance in the Secular. **c. LIWC analysis for text responses.** Out of 534 comparisons (86 textual features x two types of questions x three video clips), 17 cases of group differences were found regarding participants’ text answers; 113 cases of violation of equal of variances assumption were found, out of them 71 cases where Variance Religious > Variance Secular. **d. Cosine similarity for text responses.** We found differences in the homogeneity of the text responses only in one question towards the Kosher video, with three out of the four measures showing significant group differences.

**Fig. 2.**
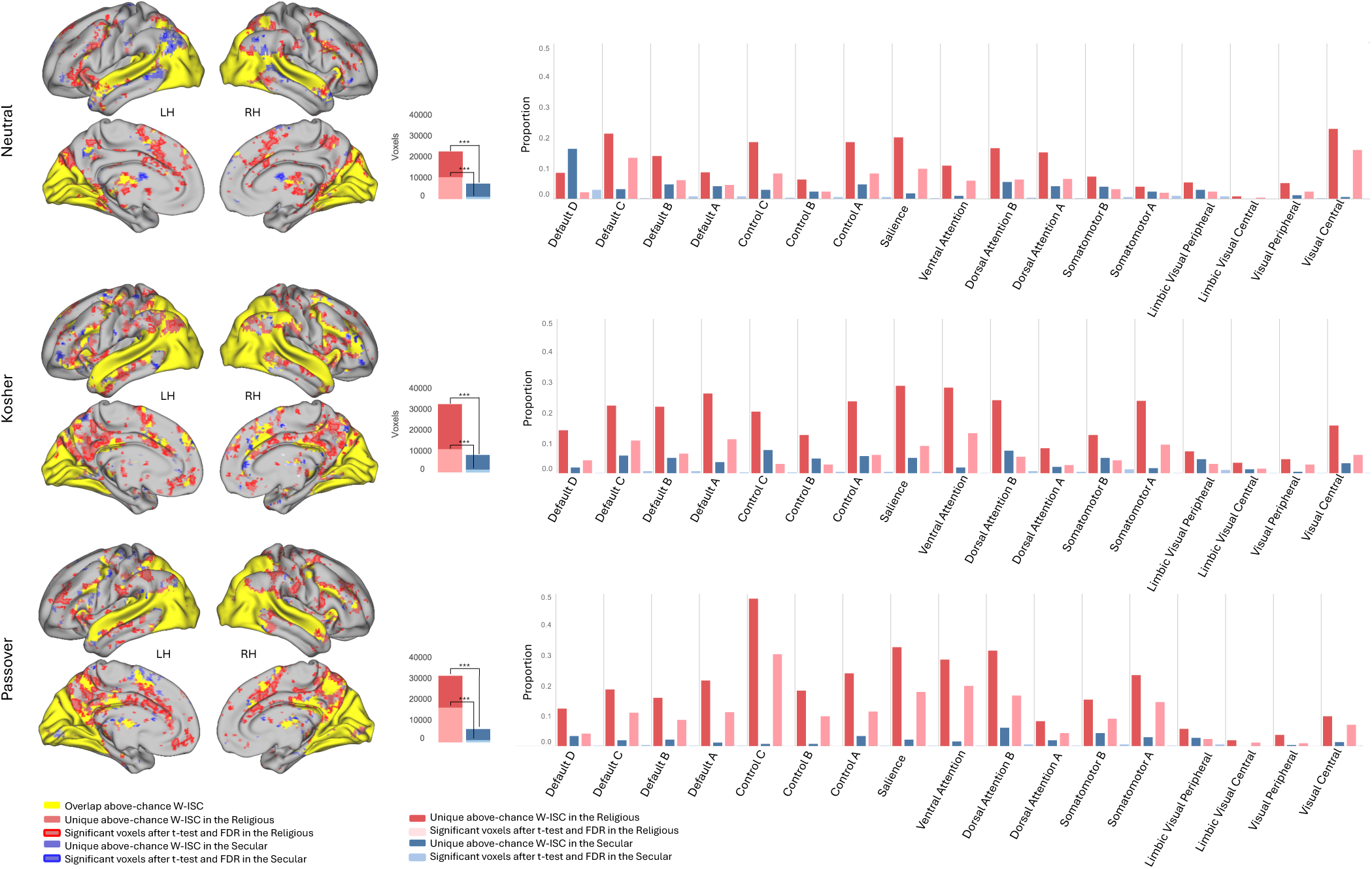
In-group neural synchronization results. **a. A comparison of the number of voxels with above-chance ISC in the Religious and Secular groups.** Red stands for Religious and blue for Secular. Full color (i.e., the full height of each column, in red / blue) represents the total number of above-chance ISC voxels. The middle portion, colored in lighter blue / red, stands for the number of unique voxels (i.e., voxels where there was an above-chance ISC score only in one group). Lighter colors mark significant voxels after a t-test and FDR correction. Note: *** *p* < .001 **b. Brain regions exhibiting inter-personal neural synchronization.** Yellow patches are for overlap regions, i.e., the ISC scores were above chance in both the Religious and Secular (mostly sensory regions); light red / blue patches are for regions where the in-group neural synchrony was unique to the Religious / Secular, respectively. Within those regions, outlined in red / blue are voxels in which the ISC differences between the groups were found significant after t-test and FDR correction (*q < .05)*. **c. Proportions of In-group Synchronized Neural Networks.** The proportions of the significant voxels in our results were computed in each of Yeo’s 17 networks. Presented are the proportions of synchronized voxels in unique voxels (i.e., voxels that had above-chance ISC score only in one group and not the other). Results present a consistent pattern of greater extent of higher in-group synchronization levels in the Religious, compared to the Secular. All the differences were found statistically significant in a Chi-squared proportion test. 65 out of the 102 the comparisons were found significant after applying Bonferroni correction for multiple comparisons.

#### Cosine Similarity

To assess whether there were group differences in in-group homogeneity of the text responses, we computed four cosine-similarity metrics: three based on TF–IDF features of tokens, lemmas, and part-of-speech tags extracted from our experimental corpus, and one based on semantic embeddings pretrained on Twitter. In five of the six videos × question comparisons, independent-samples t-tests on Fisher-z-transformed cosine scores revealed no significant group differences (p > .05). By contrast, for the Kosher video under the description question, Religious participants exhibited significantly greater in-group similarity than Secular participants on three measures. Specifically, lemma-level similarity (MReligious = 0.298; MSecular = 0.265; t(40) = –2.87, p = 0.007, d = –0.85), POS-level similarity (MReligious = 0.903; MSecular = 0.857; t(36) = –2.11, p = 0.041, d = –0.64), and semantic similarity (MReligious = 0.804; MSecular = 0.747; t(43) = –2.48, p = 0.017, d = –0.72) were all higher in the Religious group. Token-level similarity did not differ (MReligious = 0.293; MSecular = 0.273; t(38) = –1.68, p = 0.102) (Fig 1d) (full set of statistics is described in table S4 in *Supplementary Materials*).

### Higher levels of in-group neural synchronization in the Religious group

In-group neural synchronization was measured using the group’s mean In-group ISC score in each voxel (see *Methods*). To determine in which voxels the in-group synchrony level was above chance, we computed *p*-value for each voxel using a Cumulative Distribution Function (CDF), for each group and video separately.

When counting the number of voxels that exhibit above-chance synchronization, there were significantly more voxels in the Religious group compared to the Secular group, consistent across all videos. In the Neutral video, there were 58,379 voxels in the Religious group compared to 43,099 voxels in the Secular group, χ²(1, *N* = 224,821 in both groups) = 2971.38, *p* < .001. In the Kosher video, the Religious group had 85,862 voxels compared to 61,683 in the Secular group, χ²(1, *N* = 224,770 in both groups) = 5898.22, *p* < .001. Similarly, in the Passover video, the Religious group had 81,806 voxels compared to 56,427 in the Secular group, χ²(1, *N* = 224,577 in both groups) = 6731.04, *p* < .001 (Fig. 2).

To further examine the differences between the two groups, we divided the synchronized brain regions into Unique Religious/Secular (i.e., regions where high in-group synchrony was observed only in one of the groups) versus Overlap (i.e., regions in which both groups exhibited high levels of in-group synchrony). Results revealed higher levels of synchronization in the Religious group, consistent across all videos (Fig. 2b). In the Neutral video, there were 22,693 voxels that were synchronized only in the Religious group compared to 7,413 that were synchronized only in the Secular group, χ²(1, *N* = 58,379 for Religious, *N* = 43,099 for Secular) = 5580.97, *p* < .001. In the Religious group, 10,375 out of the unique voxels exhibited significant differences between the groups after the t-test and FDR correction, compared to 1,182 in the Secular group, χ²(1, *N* = 22,693 for Religious, *N* = 7,413 for Secular) = 2094.34, *p* < .001.

Similarly, in the Kosher video, the Religious group had 32,571 unique voxels compared to 8,392 in the Secular group, χ²(1, *N* = 85,862 for Religious, *N* = 61,683 for Secular) = 10,594.17, *p* < .001. After applying FDR correction, 11,001 significant unique voxels were found in the Religious group compared to 1,442 in the Secular group, χ²(1, *N* = 32,571 for Religious, *N* = 8,392 for Secular) = 868.63, *p* < .001.

The Passover video revealed 31,716 unique voxels in the Religious group compared to 6,337 in the Secular group, χ²(1, *N* = 81,806 for Religious, *N* = 56,427 for Secular) = 12,694.65, *p* < .001. After FDR correction, 16,579 significant unique voxels were observed in the Religious group compared to 1,043 in the Secular group, χ²(1, *N* = 31,716 for Religious, *N* = 6,337 for Secular) = 2724.72, *p* < .001.

In the Overlap regions a similar pattern was observed, with the Religious demonstrating higher in-group synchrony across all videos (see Fig. S3 in *Supplementary Materials*).

Next, we grouped single voxels into neural networks, based on Yeo’s division into 17 networks. We then measured the proportions of synchrony in each of the networks (Fig. 2c). This allowed us to examine in which neural networks the synchrony differed between the groups. This analysis revealed that the proportion of voxels with high synchrony level was always higher in the Religious compared to the Secular (except for one case in the Auditory network in the Neutral video). The differences were also statistically significant, as measured in Chi-squared proportion test, with 95 out of the 102 comparisons remaining significant after Bonferroni multiple comparisons. Noteworthy differences between the groups were found in the Control C network, where there were up to 64.00 times more synchronized voxels in the Religious; The Salience network, where there were up to 14.70 times more synchronized voxels in the Religious; The Ventral Attention 1 networks, where there were up to 19.35 times more synchronized voxels in the Religious; and the Default A network, where there were up to 18.22 times more synchronized voxels in the Religious (table S6 in *Supplementary Materials*).

To summarize, we found a clear and consistent pattern of higher in-group neural synchronization within the Religious group, both in the magnitude and extent.

### ISC Scores Reliably Predict Religious Affiliation

To test if it is possible to predict participants’ religious affiliation based on their level of synchronization with other members of their group, we trained an SVM classifier (see *Methods*). Our results suggest that we were able to classify participant’s affiliation in 7627 voxels in the Neutral video (accuracy between 0.6429 and 0.9195); 5653 voxels in the Kosher video (accuracy between 0.6429 and 0.8946); and 9185 in the Passover video (accuracy between 0.6429 and 0.9036). In the Neutral video, the most prominent classification voxels appeared in occipital regions (medial ventral and lateral) and the fusiform gyrus, along with the precuneus; for the Kosher video, large clusters were found both occipital regions as well as precentral gyrus, Cingulate, inferior Parietal Lobule and Precuneus; as for the Passover video, the Precuneus was most prominent, along with Cingulate gyrus and parietal lobule (Fig. 3a; see also table S7 in *Supplementary Materials*).

**Fig. 3.**
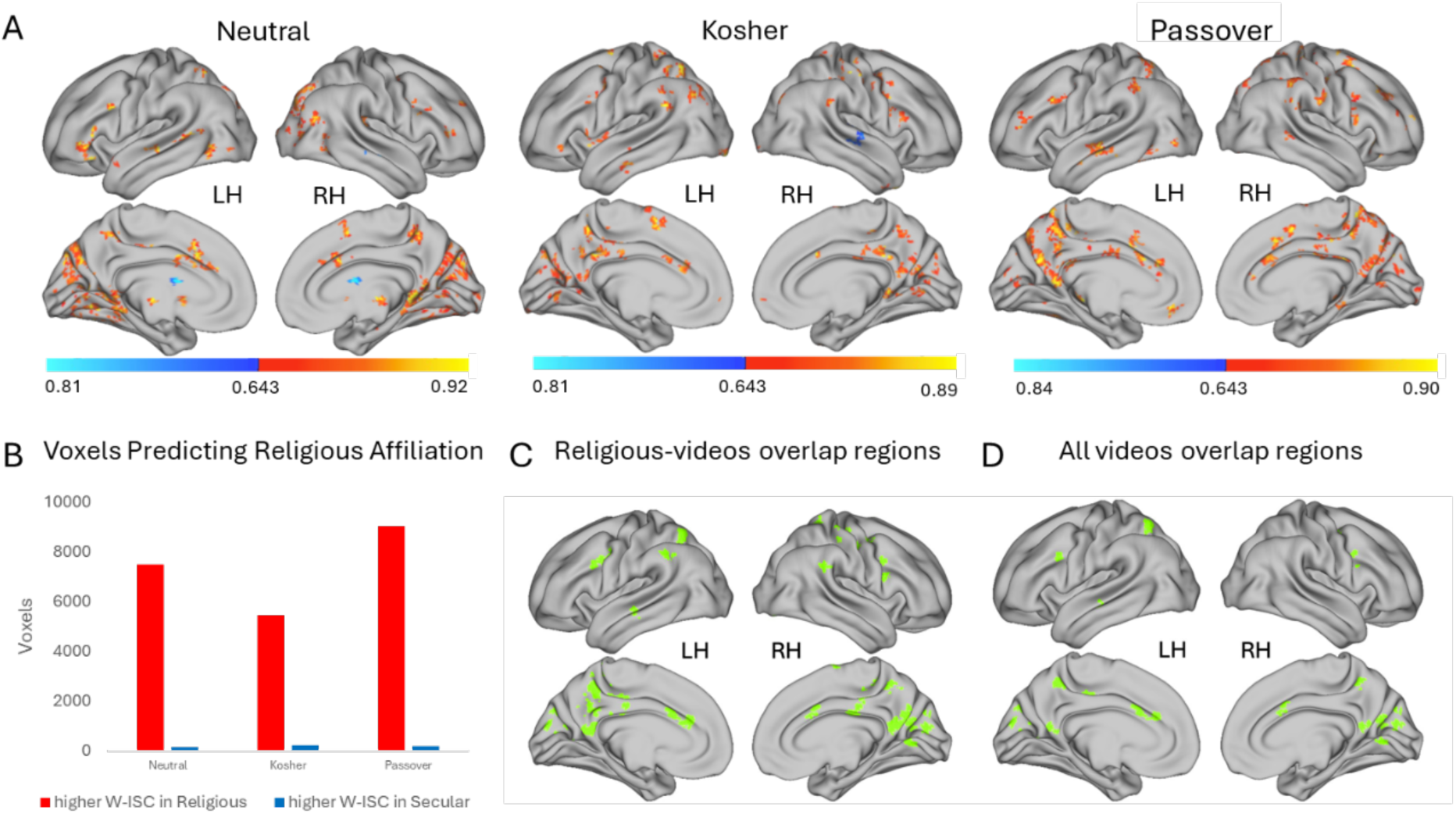
Predicting Religious Affiliation Based on ISC Scores Using an SVM Model. a. Colored in heat scale are brain regions in which an SVM model successfully predicted participants’ religious-affiliation – Religious or Secular – based on their ISC score. In warm colors are regions where the ISC scores were higher in the Religious; cool colors are for regions where the S-ISC scores were higher in the Secular. Accuracy rates presented in the heat scale. **b.** quantifying the number of voxels that predicted religious affiliation, divided by group based on higher in-group synchrony in either the Religious or Secular group. **c.** Regions in which the accuracy of the SVM classifier to predict religious affiliation – Religious or Secular, based on participants’ ISC score – was reliable across the Kosher and Passover videos. **d** Regions in which the accuracy of the SVM classifier to predict religious affiliation – Religious or Secular, based on participants’ ISC score – was reliable across all three videos.

### Cross-videos Consistency in Predicting the Religious-Affiliation in the Precuneus and Cingulate gyrus

To investigate the consistency of regions in which the SVM classifier exceeded chance, we applied a spatial expansion technique to the identified voxels in each video. We then performed two sets of intersections – across the religious-content videos (Kosher and Passover) and across all videos (see *Methods*).

The analysis revealed 4265 overlapping voxels in the religious-content videos. Upon examining the gyri with the highest concentration of these voxels, the two most prominent were the Precuneus and the Cingulate Gyrus (Fig. 3c). The number of overlapping regions decreased in the intersection across all 3 videos (including the Neutral) and resulted in 1556 voxels. As before, the gyrus that demonstrated the highest concentration of voxels was the Precuneus, followed by the Medio-Ventral Occipital Cortex and the Cingulate gyrus.

## Discussion

In this study we examined if in-group neural synchronization can be viewed as a characterizing feature of a group, testing this question in the context of religious affiliation. We compared the in-group neural synchronization of Religious vs Secular towards both religious and neutral narratives. Neuroimaging results revealed consistently higher in-group neural synchrony among Religious participants — in the magnitude and spatial extent — regardless the content. This pattern was most prominent in regions of the Control, Attention, Default and Somatomotor networks. Notably, it was possible to predict participants’ religious affiliation based on how synchronized their response in these networks was with their group members. Furthermore, this increased neural alignment was not mirrored in participants’ explicit views, as no clear pattern of higher in-group homogeneity was found within the Religious group in the behavioral measures.

Previous literature highlighted the involvement of the DMN in group-dependent neural responses, mainly towards polarizing – and not neutral – content (e.g., Leong et al., 2020; van Baar et al., 2021). However, our results revealed a consisted pattern of higher neural synchronization within the Religious group, with multiple regions, including outside the DMN, exhibiting both content specific and non-content specific increased neural synchronization.

Regions where group differences in neural synchrony were not content-specific include parts of the Control, Dorsal Attention, Salience and Default Mode networks. Neural synchronization in these high-order regions during free viewing of videos, especially in the DMN, is generally associated with shared interpretation of narratives (Hasson et al., 2008; Honey et al., 2012; Jääskeläinen et al., 2008; Kauppi et al., 2010; Ki et al., 2016; Lahnakoski et al., 2014; Lerner et al., 2011; Yeshurun et al., 2017) and shared emotional processing (Bacha-Trams et al., 2017; Finn et al., 2018; Nummenmaa et al., 2012; van Baar et al., 2021), where distinct neural responses in these regions serve as a marker for individuals’ socio-cultural group (Andrews et al., 2019; Hakonen et al., 2022; Leong et al., 2020). Regions that demonstrated a content-specific pattern, i.e., differences in in-group neural synchronization were primarily towards religious narratives, include mostly parts of the Ventral Attention network, and interestingly, Somatomotor regions.

Neural synchrony in Somatomotor regions was found to be correlated with high emotional arousal (Nummenmaa et al., 2012) and shared mental action simulation (Nummenmaa et al., 2014). It was also found in right-wing individuals while they were watching right-wing political video clips (Katabi et al., 2023). The latter study raised the hypothesis that individuals use sensorimotor simulative representation to process engaging narratives, which might facilitate their ability to understand the intentions of social agents’ actions and identify with them. This hypothesis is further supported by our current findings, given the content-specific synchronization pattern.

Interestingly, high in-group neural synchronization in the Religious group was found also in the Salience network (that is composed mainly of the anterior Insula (AI) and dorsal anterior Cingulate cortex (dACC) (Menon, 2015)). This network encodes differentiation of group identities based on social categories, responds to group affiliations saliency and process emotional significance of in- vs out-group distinctions (Cikara et al., 2017). Group salience also shifts self-concept from “I” to “we”, altering the neural responses in the DMN, that supports social identity representation (Cikara & Van Bavel, 2014). Thus, synchronized neural responses in the Salience network (along the DMN) might strength group identification and potentially facilitate “us vs them” perspective.

Importantly, the overall higher in-group neural synchronization of the Religious group cannot be attributed to semantic usage of familiar religious terminology; only one video (Kosher) presented a religious perspective and concepts, whereas the neural synchronization pattern was consistent across all three video-clips. Additionally, although engagement with a narrative synchronizes neural responses (Ohad & Yeshurun, 2023), Religious’ neural synchronization cannot be attributed solely to that since group-differences in engagement were found only towards one video (Kosher). Furthermore, the variances of the Religious group members’ scores in the explicit measures (agreement, engagement and emotional responses towards the videos) did not significantly differ from those of the Secular. Similarly, in two text analysis approaches we applied on participants’ responses, no group differences were found in terms of homogeneity. This dissociation between neuroimaging and explicit measures implies that neural synchrony might capture a deeper, more complex shared representation across group members, exceeding sheer self-report interpretation and attitudes.

Past studies that examined group-differences in neural synchronization primarily focused on which brain regions exhibit group-dependent neural responses, showing that group differences in interpretation (either experimentally manipulated differences or those who stem from having different backgrounds) are reflected in distinct neural responses (e.g., Bacha-Trams et al., 2020; Hakonen et al., 2022; Katabi et al., 2023; Lahnakoski et al., 2014; Leong et al., 2020; Yeshurun et al., 2017; Zadbood et al., 2022). Increased neural synchronization across individuals was viewed as a phenomenon related to cognitive and interpersonal functions, such as successful communication (Stephens et al., 2010) and having a more similar interpretation of narratives (Nguyen et al., 2019). Yet, the prominent and consistent pattern of the increased neural synchronization within the Religious groups, prevailing across all neural networks and stimuli, implies that the degree of in-group neural synchronization acts as a potential biomarker for groups’ social structure. Not only we found significantly larger proportions of networks exhibiting higher ingroup neural synchronization in the Religious groups, it was also possible to predict individuals’ religious affiliation solely based on their in-group neural synchrony (with accuracy scores of up to 92%), as shown in our SVM classification results, demonstrating the robustness of the group differences in neural synchronization.

We suggest that the higher in-group neural synchronization of the religious individuals can be explained by a more homogenous structure of their group. Religious societies are characterized by higher degree of collectivism, which fosters group cohesion (Abrams & Hogg, 1990) and religious individuals are inclined to endorse moral values that serve group cohesion (Ståhl, 2021). These attributes, together with shared socialization processes, drive a more homogenous social structure (Ruiter & van Tubergen, 2009). In Israel there is a separate religious education system, that along with effective socialization mechanisms, contributes to greater unity of the religious society (Katsman, 2020; Maoz, 2005; Shamai, 2000). These factors are prone to result in greater group homophily, which promotes closer social distances within its members (McPherson et al., 2001). It was shown that neural responses during free viewing movie clips were more similar among individuals who were closer to each other in their shared social networks (Parkinson et al., 2018). We provide compelling evidence demonstrating this principle on a broader scale, with religious individuals exhibiting increased neural synchronization as a possible reflection of higher group cohesion.

Along with a more homogenous structure, we propose that potential group differences in cognitive styles might contribute to increased neural synchronization among religious individuals. It was previously found that individuals with a greater need for certainty exhibit higher tendency for in-group neural synchronization (van Baar et al., 2021). Although we did not measure uncertainty-intolerance in the current study, religious belief systems address and fulfil the need for certainty, doing so through strict rules and rituals, which reduce cognitive flexibility (Zmigrod et al., 2019). More rigid cognitive style, shaped through repetitive religious rituals, might serve as a mechanism that enhances neural synchronization within religious societies.

While our findings provide valuable insights regarding social groups neural synchronization patterns, several limitations should be acknowledged. First, we consider two aspects in the experimental design that might have compromised our behavioral measurements. During the fMRI scan we asked participants to rate their agreement level towards the video; yet, the Passover video clip presents conflicting opinions, so it might have been confusing answering about agreement. Second, participants answered the post scan questionnaire, with questions about their emotional response towards the videos, approximately an hour after they watched the videos. It is reasonable that the intensity of their emotions decreased, so their scores did not capture their real-time emotional response. Thus, further group differences in participants’ explicit interpretations of the narratives may have been obscured.

We propose that by taking additional behavioral measures on cognitive and social traits, future studies could offer a more elaborate mechanism to explain the higher in-group neural synchronization among the religious participants. For example, measuring intolerance of certainty (van Baar et al., 2021), loneliness (Baek et al., 2023), cognitive inflexibility (Zmigrod, 2020; Zmigrod et al., 2019) and in-group favoritism (Johnson et al., 2012) which might explain some of the increased synchronization in our results.

To our knowledge, this study is the first to test the relation between neural synchronization and religious affiliation. We found robust evidence to support the notion of increased in-group neural synchronization among the Religious group, which exceeds the religious content and was evident for neutral content as well. We demonstrated that this pattern is prominent not only in the DMN, but also in Control, Dorsal Attention, Salience, and Somatomotor networks. Taken together, our findings suggest that in-group neural synchronization might be a marker for social groups’ structure, and serve as a first step in enabling a deeper understanding of group dynamic mechanisms that drive shared neural responses among individuals.

## Conflict of interest statement

The authors declare no competing financial interests.

## Acknowledgments

We thank Eldad Aviv for his help with the text data analysis and Hadar Nakar for her assistance in the MRI scanning. This work was supported by the Israel Science Foundation Grant No. 388/24.

## Data & Code Availability

All behavioral data and preprocessed neuroimaging data are available in a public Open Science Framework (OSF) repository [http://tiny.cc/OSF_repo]. The code that was used for the statistical analysis is available in a public GitHub repository [https://github.com/YohayZvi/religion-neuro-sync].

## Notes

http://tiny.cc/OSF_repo

https://github.com/YohayZvi/religion-neuro-sync

### Competing Interest Statement

The authors have declared no competing interest.

