## Supplementary material for "You Are What You Sync: Neural Synchronization Predicts Religious Affiliation": Supplamental Materials

**Supplemental Materials**

**Preliminary Screening Questionnaire**

1. Religious affiliation:
    a. Atheist
    b. Jew
    c. Chrisitan
    d. Muslim
    e. Other: ______
2. Are you:
    a. Secular
    b. Religious
    c. Ba'al Teshuva (converted into religious)
    d. Ex-religious
3. How would you define the level of religiosity in the home where you grew-up?
    a. Secular
    b. Traditional
    c. Conservative
    d. Reform Judaism
    e. National-religious
    f. Ultra-orthodox (Haredi')
    g. Other: ______
4. How would you define your level of religiosity?
    a. Religious
    b. National religious
    c. Reform Jew
    d. Conservative
    e. Traditional
    f. Secular
    g. Other: ______
5. Are you an Ex-religious?
    a. No
    b. Yes
        i. At what age / range of ages did you become non-religious? _________
        ii. Was the process of moving away from religion done:
            1. Alone
            2. With some of the family members
            3. With the entire family
        iii. How many years have passed since you became non-religious?
6. Are you Ba'al Teshuva (converted into religious)?
    a. Yes

    b. No
        i. At what age (or age range) did you become religious? _________
        ii. Was the process of turning into religious done:
            1. Alone
            2. With some of the family members
            3. With the entire family
        iii. How many years have passed since you became religious?

7. Please choose the circles combination that symbolizes best your group affiliation

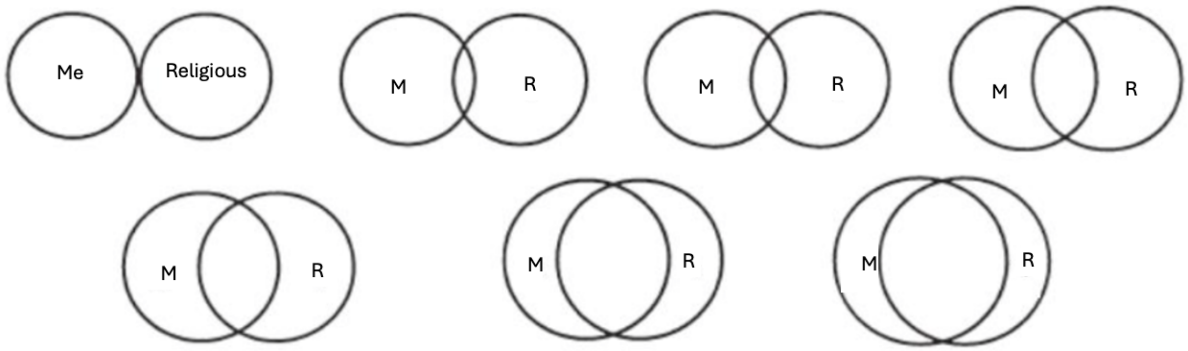


8. Have you gone under more than one religious “transition” in your lifetime? (for example: been secular, turned into religious, and became secular again):
    a. Yes
    b. No
    c. Other: _________
9. How many of your close friends and acquaintances are religious?
    a. All my friends and acquaintances
    b. Many of my friends and acquaintances
    c. Some of my friends and acquaintances
    d. Very few of my friends and acquaintances
    e. None of my friends and acquaintances
10. Gender:
    a. Male
    b. Female
    c. Other: _________
11. Proficiency in spoken and comprehension skills in Hebrew:
    a. Native speaker
    b. Near-native fluency
    c. Advanced
    d. Intermediate
    e. Beginner
12. Age: _________
13. First name: _________
14. Email address (In order to contact you in case you will be found fit for the study): _________

### Post-scan Questionnaire

**The Weather video-clip\ The Chametz in Hospitals video-clip\ The Alternative Kosher video-clip**

Describe shortly what you saw in the video:

___________________________________

What is the first thing that pops to your head after watching the video?

___________________________________

Do you support or against the stand that was presented in the video?

1) Support
2) Against
3) Other, please elaborate: ______________

How much does the video make you feel:

| | Not at all | A little | Moderately | A lot | Very much |
|---|---|---|---|---|---|
| Happiness | | | | | |
| Sadness | | | | | |
| Fear | | | | | |
| Disgust | | | | | |
| Anger | | | | | |
| Surprise | | | | | |

Have you watched this video before?

1) Yes
2) No
3) I don't remember

Are you (please choose the most accurate answer for you):

1) Secular
2) Religious
3) Ex-religious
4) Repent

How would you define your current level of religiosity?

1) Secular
2) Traditional

3) National religious
4) Ultra-orthodox ('Haredi')
5) Other: ________

How many of your friends and acquaintances are religious?

1) All my friends and acquaintances
2) Many of my friends and acquaintances
3) Very few of my friends and acquaintances
4) None of my friends and acquaintances

Gender:

1) Male
2) Female
3) Other

Age: ______

What do you think is the purpose of the study?

______________________________________

Do you have any comments about the entire experiment?

______________________________________

**Null distributions statistics**

*Table S1a – ISC null distributions' statistical parameters.*

| Group | Video | Voxels | Mean | Median | SD | Min | Q1 | Q2 | Q3 | Max |
|---|---|---|---|---|---|---|---|---|---|---|
| Religious | Neutral | 224821 | 0.000 | 0.000 | 0.037 | -0.479 | -0.020 | 0.000 | 0.020 | 0.458 |
| Secular | Neutral | 224821 | 0.000 | 0.000 | 0.034 | -0.441 | -0.019 | 0.000 | 0.019 | 0.443 |
| Religious | Kosher | 224770 | -0.001 | -0.001 | 0.034 | -0.467 | -0.019 | -0.001 | 0.018 | 0.451 |
| Secular | Kosher | 224770 | 0.000 | 0.000 | 0.033 | -0.452 | -0.018 | 0.000 | 0.018 | 0.464 |
| Religious | Passover | 224577 | 0.000 | 0.000 | 0.046 | -0.637 | -0.019 | 0.000 | 0.019 | 0.604 |
| Secular | Passover | 224577 | 0.000 | 0.000 | 0.043 | -0.642 | -0.018 | 0.000 | 0.018 | 0.631 |

*Table S1b – SVM accuracy rates null distributions' statistical parameters. The 95$^{th}$ percentile was the accuracy rate threshold.*

| Video | Voxels | Mean | Median | SD | Min | Q1 | Q2 | Q3 | 95th percentile |
|---|---|---|---|---|---|---|---|---|---|
| Neutral | 65792 | 0.5483 | 0.5476 | 0.0569 | 0.2857 | 0.5000 | 0.5476 | 0.5952 | 0.6429 |
| Kosher | 94254 | 0.5477 | 0.5476 | 0.0569 | 0.2619 | 0.5000 | 0.5476 | 0.5952 | 0.6429 |
| Passover | 88143 | 0.5482 | 0.5476 | 0.0564 | 0.2857 | 0.5000 | 0.5476 | 0.5952 | 0.6429 |

**Tuning grid results for the clusters size correction**

*Fig. S1 – Tuning grid results of the SVM cluster size correction*

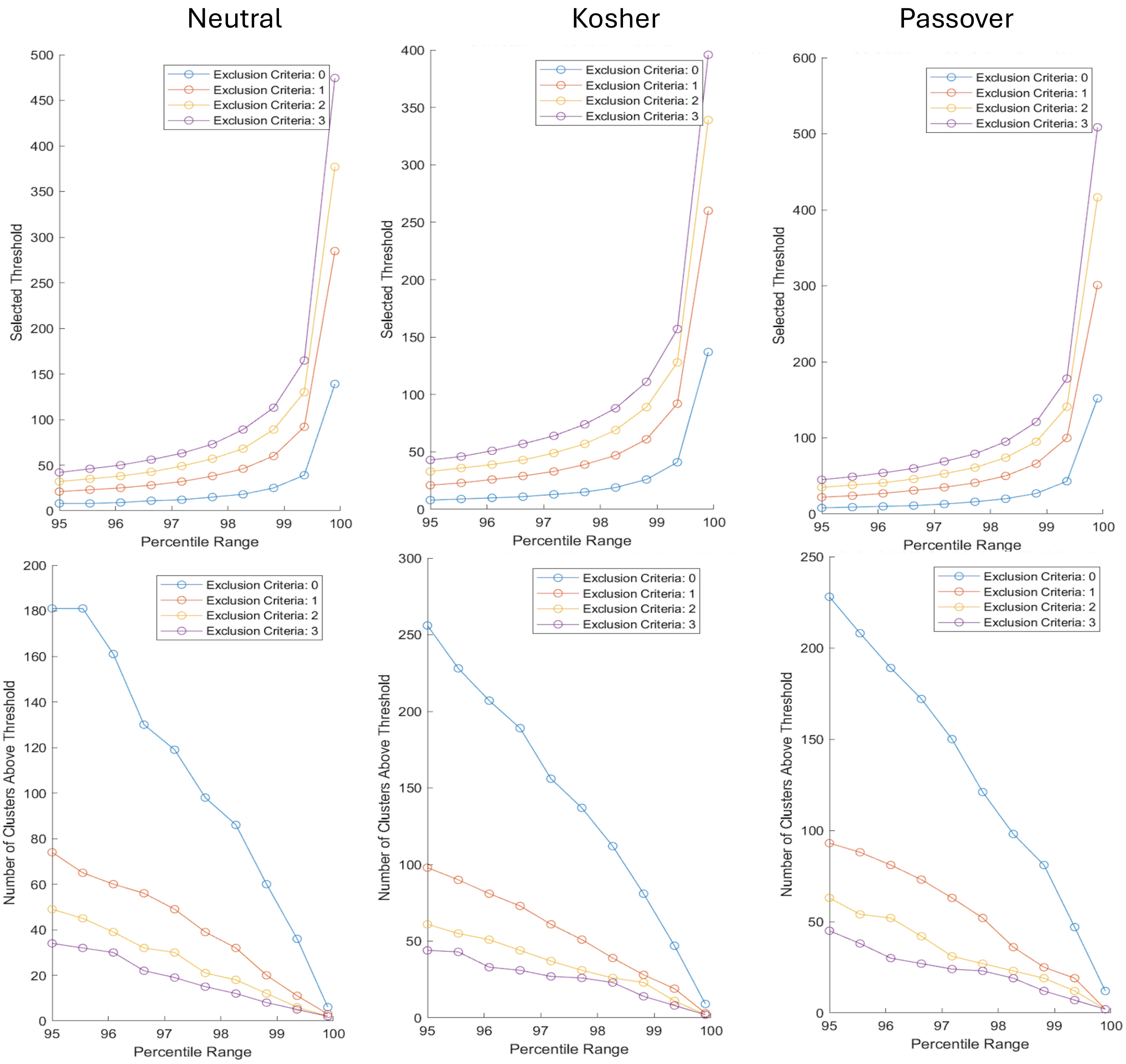


We used a tuning grid of clusters' size exclusion criteria (0,1,2,3) and linear space of 10 values between 95 to 99.9 as percentiles of the clusters size in the null distribution. Chosen threshold was the 99.356 percentile with no exclusion of cluster size.

**Agreement and engagement scores during the scan**

*Table S2a – Mann-Whitney U test results for the agreement and engagement scores during the scan.*

| | U | df | p |
|---|---|---|---|
| Neutral agreement | 242.500 | | 0.250 |
| Neutral interest | 223.500 | | 0.524 |
| Neutral emotional Involved | 161.000 | | 0.303 |
| Passover agreement | 218.000 | | 0.626 |
| Passover interest | 267.000 | | 0.070 |
| Passover emotional Involved | 264.500 | | 0.081 |
| Kosher agreement | 319.000 | | 0.001 |
| Kosher interest | 289.000 | | 0.016 |
| Kosher emotional Involved | 311.500 | | 0.003 |

*Note*. Mann-Whitney U test.

*Table S2b – Test of Equality of Variances (Levene's) for the agreement and engagement questions.*

| | F | $df_1$ | $df_2$ | p |
|---|---|---|---|---|
| Neutral agreement | 0.358 | 1 | 38 | 0.553 |
| Neutral interest | 0.052 | 1 | 38 | 0.820 |
| Neutral emotional Involved | 0.001 | 1 | 38 | 0.973 |
| Passover agreement | 0.472 | 1 | 38 | 0.496 |
| Passover interest | 0.003 | 1 | 38 | 0.956 |
| Passover emotional Involved | 0.831 | 1 | 38 | 0.368 |
| Kosher agreement | 2.918 | 1 | 38 | 0.096 |
| Kosher interest | 8.904 | 1 | 38 | 0.005 |
| Kosher emotional Involved | 3.604 | 1 | 38 | 0.065 |

*Table S2c – Descriptives of the agreement and engagement scores.*

| | Group | N | Mean | SD | SE | Coefficient of variation | Mean Rank | Sum Rank |
|---|---|---|---|---|---|---|---|---|
| Neutral agreement | Religious | 19 | 68.452 | 17.377 | 3.987 | 0.254 | 22.763 | 432.500 |
| | Secular | 21 | 63.714 | 15.942 | 3.479 | 0.250 | 18.452 | 387.500 |
| Neutral interest | Religious | 19 | 58.265 | 24.977 | 5.730 | 0.429 | 21.763 | 413.500 |
| | Secular | 21 | 52.406 | 23.332 | 5.091 | 0.445 | 19.357 | 406.500 |
| Neutral emotional Involved | Religious | 19 | 34.536 | 21.034 | 4.826 | 0.609 | 18.474 | 351.000 |
| | Secular | 21 | 39.503 | 18.569 | 4.052 | 0.470 | 22.333 | 469.000 |
| Passover agreement | Religious | 19 | 59.655 | 24.636 | 5.652 | 0.413 | 21.474 | 408.000 |
| | Secular | 21 | 55.457 | 28.560 | 6.232 | 0.515 | 19.619 | 412.000 |
| Passover interest | Religious | 19 | 75.428 | 14.729 | 3.379 | 0.195 | 24.053 | 457.000 |
| | Secular | 21 | 67.402 | 14.189 | 3.096 | 0.211 | 17.286 | 363.000 |
| Passover emotional Involved | Religious | 19 | 73.025 | 18.578 | 4.262 | 0.254 | 23.921 | 454.500 |
| | Secular | 21 | 61.467 | 22.103 | 4.823 | 0.360 | 17.405 | 365.500 |
| Kosher agreement | Religious | 19 | 47.499 | 32.003 | 7.342 | 0.674 | 26.789 | 509.000 |
| | Secular | 21 | 17.600 | 25.549 | 5.575 | 1.452 | 14.810 | 311.000 |
| Kosher interest | Religious | 19 | 72.821 | 15.803 | 3.625 | 0.217 | 25.211 | 479.000 |
| | Secular | 21 | 48.968 | 28.324 | 6.181 | 0.578 | 16.238 | 341.000 |
| Kosher emotional Involved | Religious | 19 | 77.518 | 15.117 | 3.468 | 0.195 | 26.395 | 501.500 |
| | Secular | 21 | 52.779 | 27.134 | 5.921 | 0.514 | 15.167 | 318.500 |

*Table S3a - Mann-Whitney U test results for post scan emotional response questionnaire.*

*Independent Samples T-Test*

| | U | df | p |
|---|---|---|---|
| Neutral-happy | 245.500 | | 0.476 |
| Neutral-sad | 206.500 | | 0.723 |
| Neutral-fear | 173.000 | | 0.195 |
| Neutral-disgust | 210.000 | | 0.573 |
| Neutral-anger | 184.500 | | 0.213 |
| Neutral-surprise | 200.500 | | 0.582 |
| Passover-happy | NaN[a] | | |
| Passover-sad | 197.500 | | 0.560 |
| Passover-fear | 180.000 | | 0.227 |
| Passover-disgust | 173.500 | | 0.202 |
| Passover-anger | 91.000 | | < .001 |
| Passover-surprise | 234.500 | | 0.695 |
| Kosher-happy | NaN[b] | | |
| Kosher-sad | 251.500 | | 0.421 |
| Kosher-fear | 261.000 | | 0.248 |
| Kosher-disgust | 217.000 | | 0.935 |
| Kosher-anger | 193.500 | | 0.493 |
| Kosher-surprise | 256.000 | | 0.320 |

*Note.* Mann-Whitney U test.

[a] The variance in Passover-happy is equal to 0 after grouping on Religious Affiliation

[b] The variance in Kosher-happy is equal to 0 after grouping on Religious Affiliation

*Table S3b - Descriptives for post scan emotional questionnaire scores.*

| question | Group | Valid | Missing | Median | Mean | Std | IQR | Min. | Max. |
|---|---|---|---|---|---|---|---|---|---|
| Neutral-happy | religious | 21 | 0 | 1 | 1.667 | 0.856 | 1 | 1 | 3 |
| Neutral-happy | secular | 21 | 0 | 1 | 1.476 | 0.75 | 1 | 1 | 3 |
| Neutral-sad | religious | 21 | 0 | 2 | 2.095 | 0.889 | 2 | 1 | 4 |
| Neutral-sad | secular | 21 | 0 | 2 | 2.238 | 1.091 | 2 | 1 | 4 |
| Neutral-fear | religious | 21 | 0 | 1 | 1.619 | 0.973 | 1 | 1 | 4 |
| Neutral-fear | secular | 21 | 0 | 2 | 1.952 | 1.117 | 1 | 1 | 5 |
| Neutral-disgust | religious | 21 | 0 | 1 | 1.095 | 0.436 | 0 | 1 | 3 |
| Neutral-disgust | secular | 21 | 0 | 1 | 1.19 | 0.68 | 0 | 1 | 4 |
| Neutral-anger | religious | 21 | 0 | 1 | 1.19 | 0.512 | 0 | 1 | 3 |
| Neutral-anger | secular | 21 | 0 | 1 | 1.619 | 1.117 | 1 | 1 | 4 |
| Neutral-surprise | religious | 21 | 0 | 1 | 1.619 | 1.024 | 1 | 1 | 5 |
| Neutral-surprise | secular | 21 | 0 | 1 | 1.667 | 0.796 | 1 | 1 | 3 |
| Passover-happy | religious | 21 | 0 | 1 | 1.429 | 0.811 | 1 | 1 | 4 |
| Passover-happy | secular | 21 | 0 | 1 | 1 | 0 | 0 | 1 | 1 |
| Passover-sad | religious | 21 | 0 | 2 | 2.714 | 1.309 | 2 | 1 | 5 |
| Passover-sad | secular | 21 | 0 | 3 | 2.905 | 1.091 | 2 | 1 | 5 |
| Passover-fear | religious | 21 | 0 | 1 | 1.429 | 0.926 | 0 | 1 | 4 |
| Passover-fear | secular | 21 | 0 | 1 | 1.714 | 1.102 | 1 | 1 | 5 |
| Passover-disgust | religious | 21 | 0 | 1 | 1.667 | 1.017 | 1 | 1 | 4 |
| Passover-disgust | secular | 21 | 0 | 2 | 2.095 | 1.179 | 2 | 1 | 4 |
| Passover-anger | religious | 21 | 0 | 2 | 2.238 | 1.136 | 2 | 1 | 5 |
| Passover-anger | secular | 21 | 0 | 4 | 3.524 | 1.03 | 1 | 1 | 5 |
| Passover-surprise | religious | 21 | 0 | 1 | 1.524 | 0.68 | 1 | 1 | 3 |
| Passover-surprise | secular | 21 | 0 | 1 | 1.429 | 0.598 | 1 | 1 | 3 |
| Kosher-happy | religious | 21 | 0 | 1 | 1.286 | 0.561 | 0 | 1 | 3 |
| Kosher-happy | secular | 21 | 0 | 1 | 1 | 0 | 0 | 1 | 1 |
| Kosher-sad | religious | 21 | 0 | 2 | 2.286 | 1.231 | 2 | 1 | 4 |
| Kosher-sad | secular | 21 | 0 | 2 | 1.952 | 1.071 | 1 | 1 | 5 |
| Kosher-fear | religious | 21 | 0 | 1 | 1.905 | 1.136 | 2 | 1 | 4 |
| Kosher-fear | secular | 21 | 0 | 1 | 1.619 | 1.244 | 1 | 1 | 5 |
| Kosher-disgust | religious | 21 | 0 | 2 | 1.905 | 1.179 | 1 | 1 | 5 |
| Kosher-disgust | secular | 21 | 0 | 1 | 2 | 1.265 | 2 | 1 | 5 |
| Kosher-anger | religious | 21 | 0 | 3 | 2.476 | 1.209 | 2 | 1 | 5 |
| Kosher-anger | secular | 21 | 0 | 3 | 2.762 | 1.446 | 3 | 1 | 5 |
| Kosher-surprise | religious | 21 | 0 | 2 | 1.714 | 0.902 | 1 | 1 | 4 |
| Kosher-surprise | secular | 21 | 0 | 1 | 1.476 | 0.75 | 1 | 1 | 3 |

### **Text analysis results**

*LIWC results*

LIWC's output includes 89 features for each individual text response, with score between 0 and 100 for each feature. We produced this output for all three video-clips (Neutral, Kosher, Passover) for each type of question (association and description). For each feature, in each question and video, we conducted t-test between the Religious and Secular. We found a total of 17 group differences (out of 534 comparisons) with no consistency, i.e., no group differences were found more than once in a single feature.

For descriptive purposes, we visualized the results only for features that at least 20% of the responses had non-zero scores.

Note: * marks group differences ($p < 0.05$); x marks violation of the equal of variances assumption ($p < 0.05$).

*Fig. S2a – LIWC results for the description question, Neutral video*

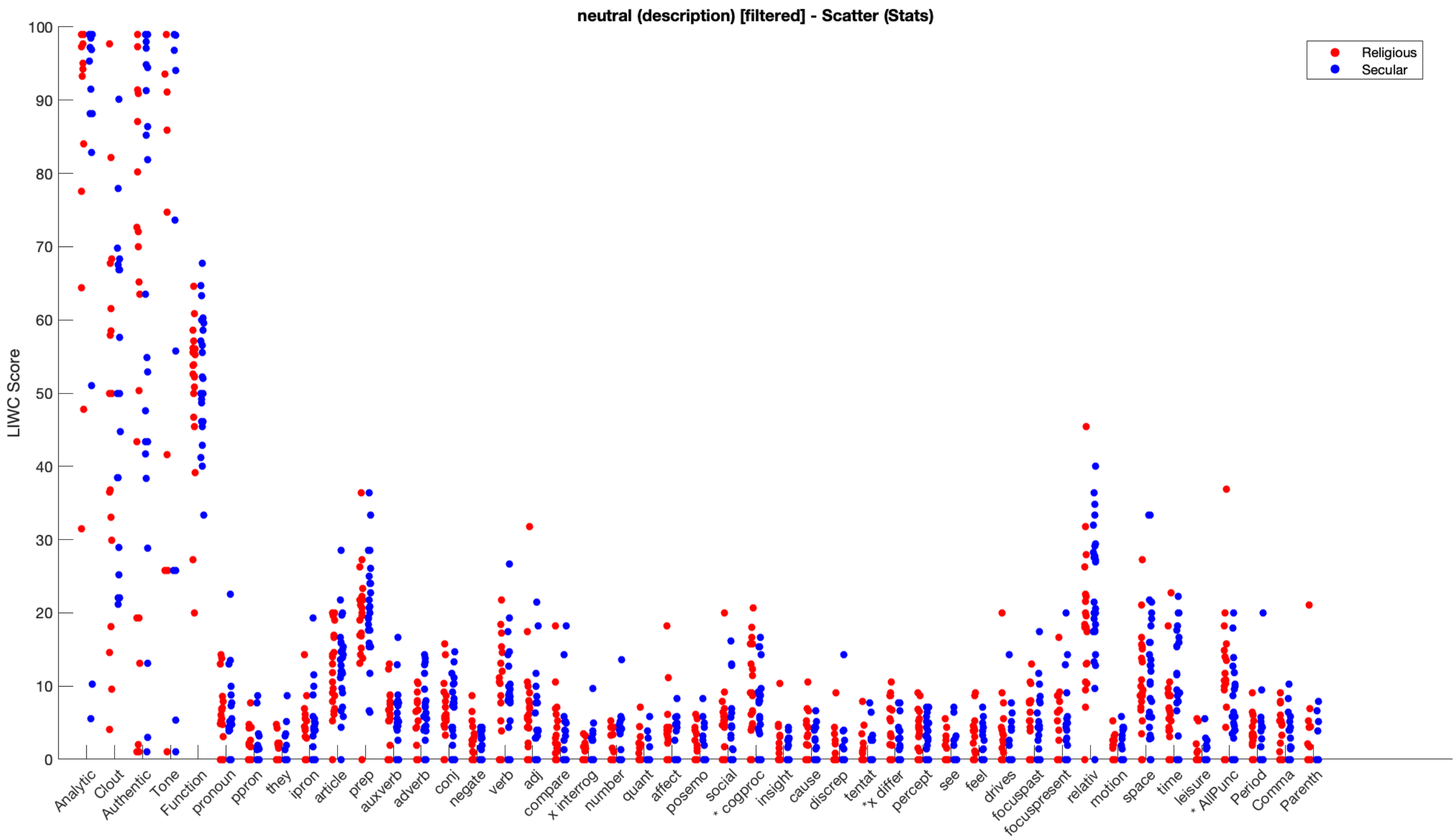


*Fig. S2b – LIWC results for the association question, Neutral video*

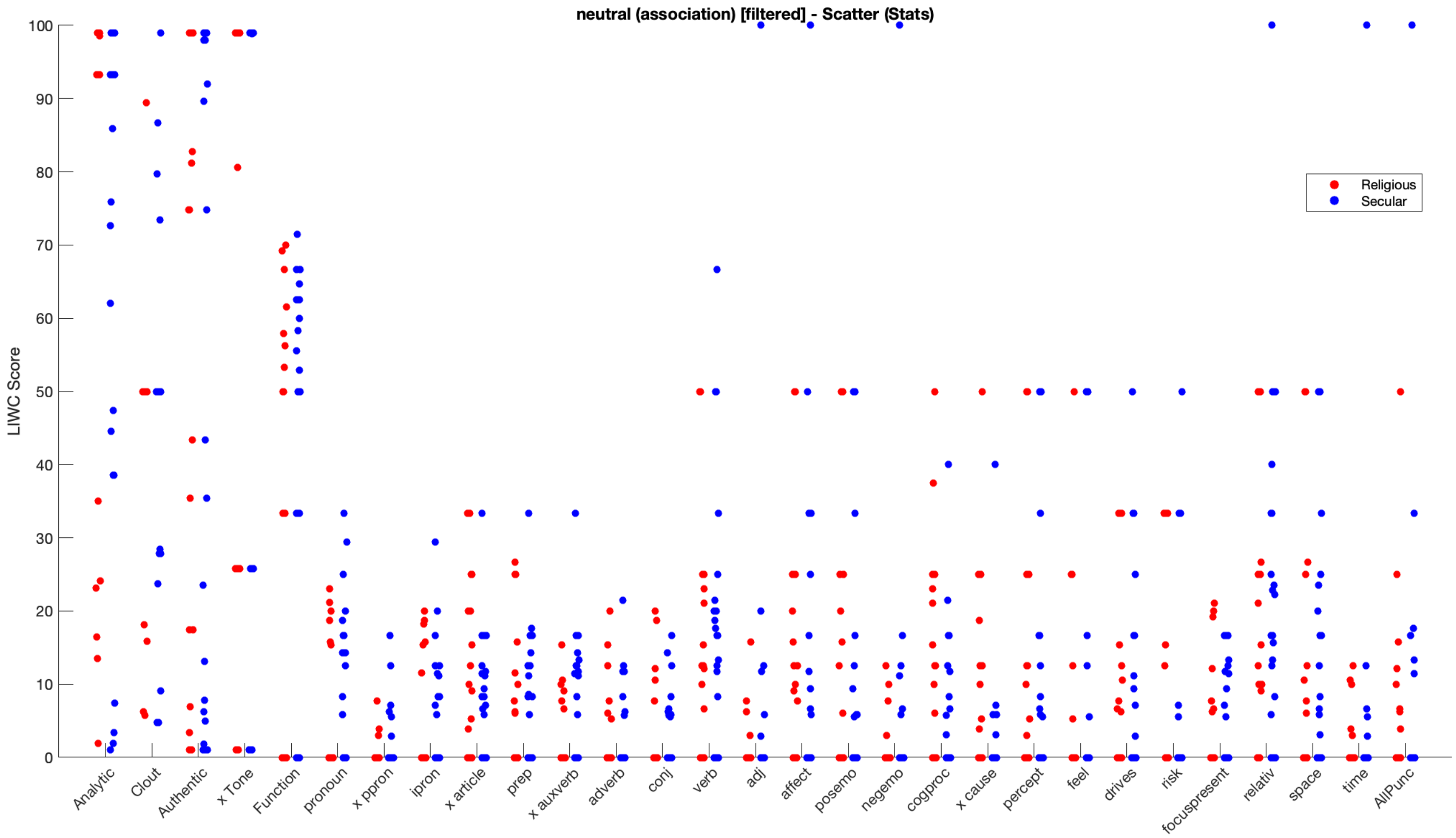

*Fig. S2c – LIWC results for the description question, Kosher video*

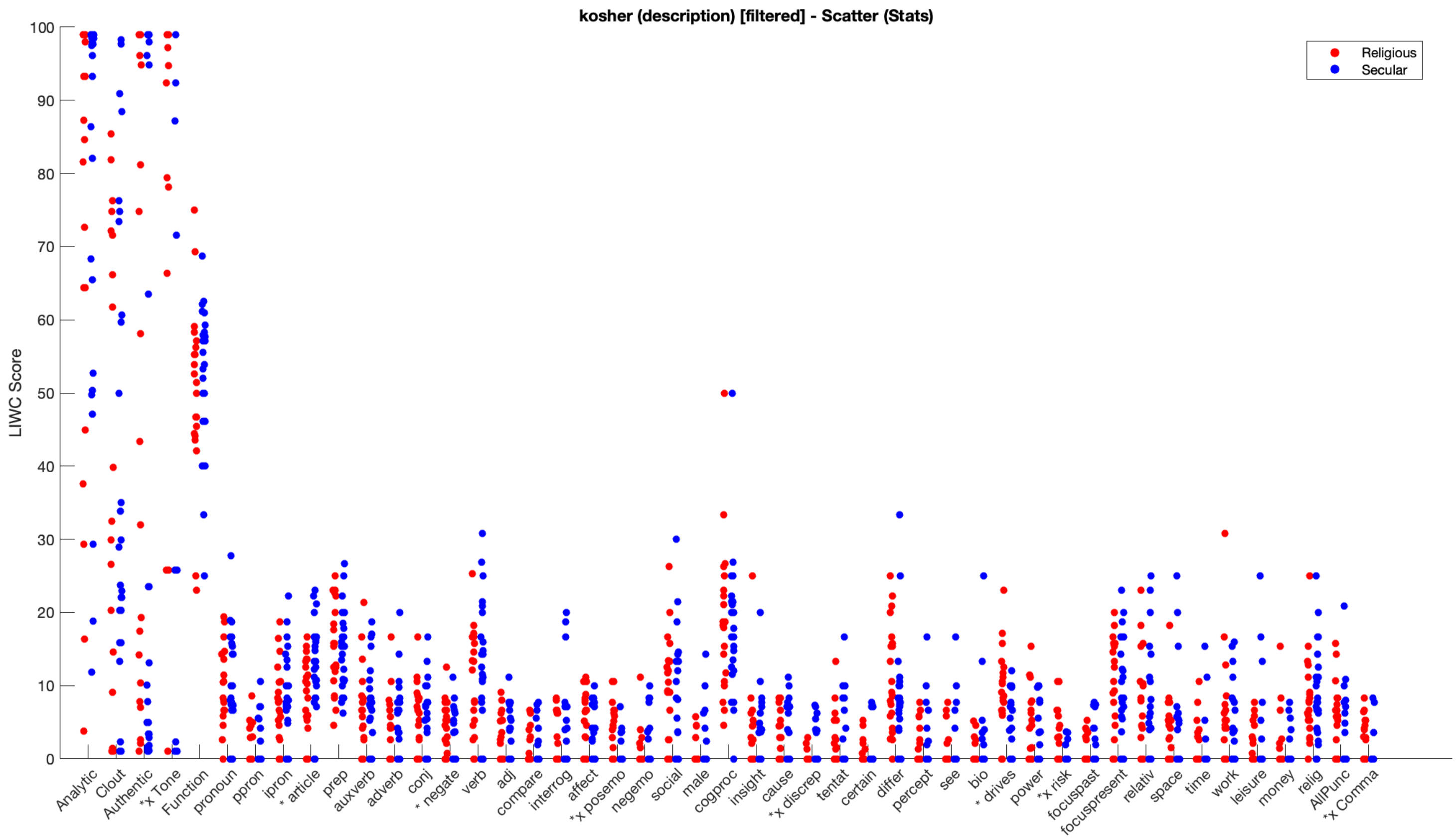


*Supp Fig. 2d – LIWC results for the association question, Kosher video*

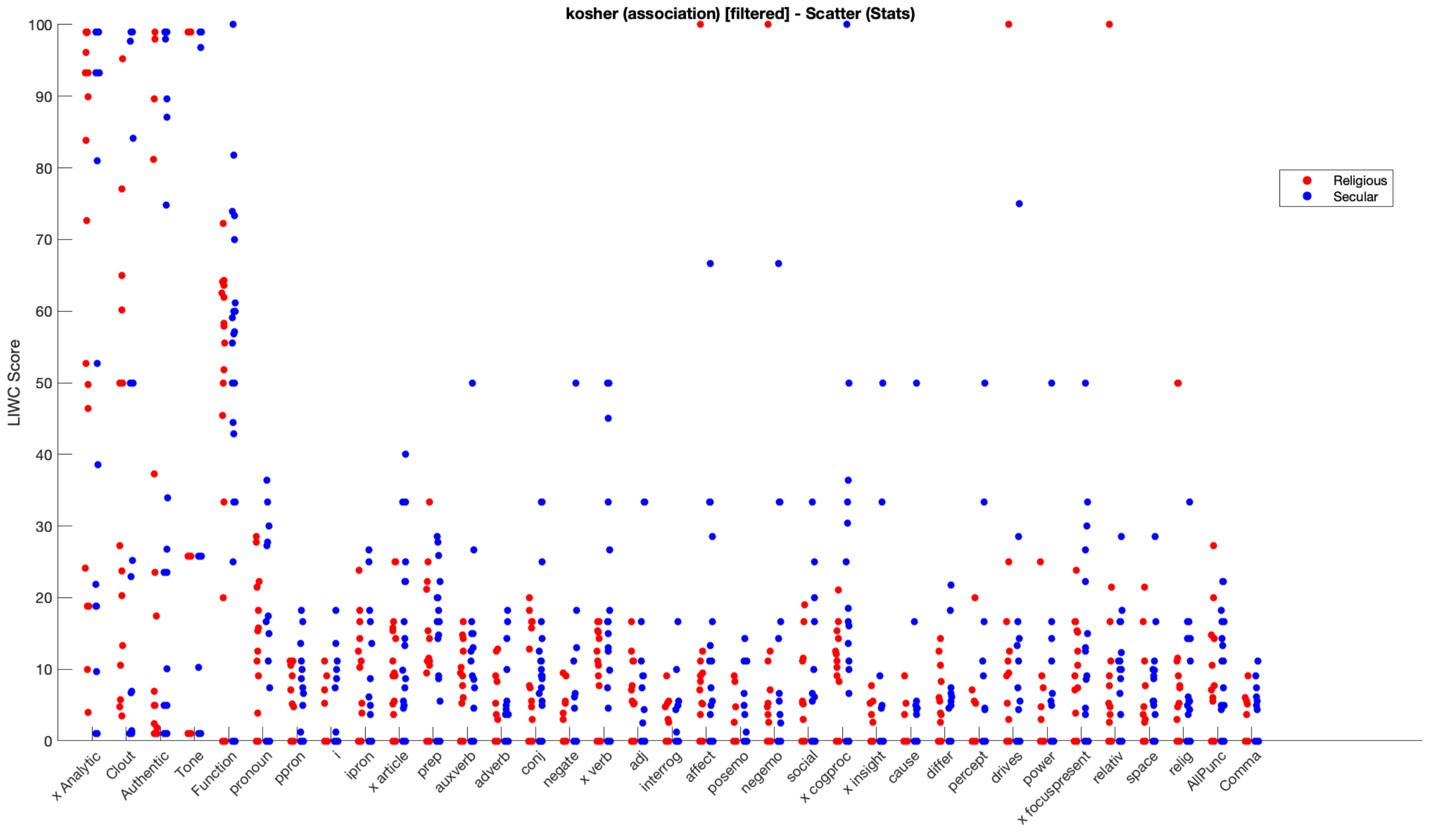

*Supp Fig. 2e – LIWC results for the description question, Passover video*

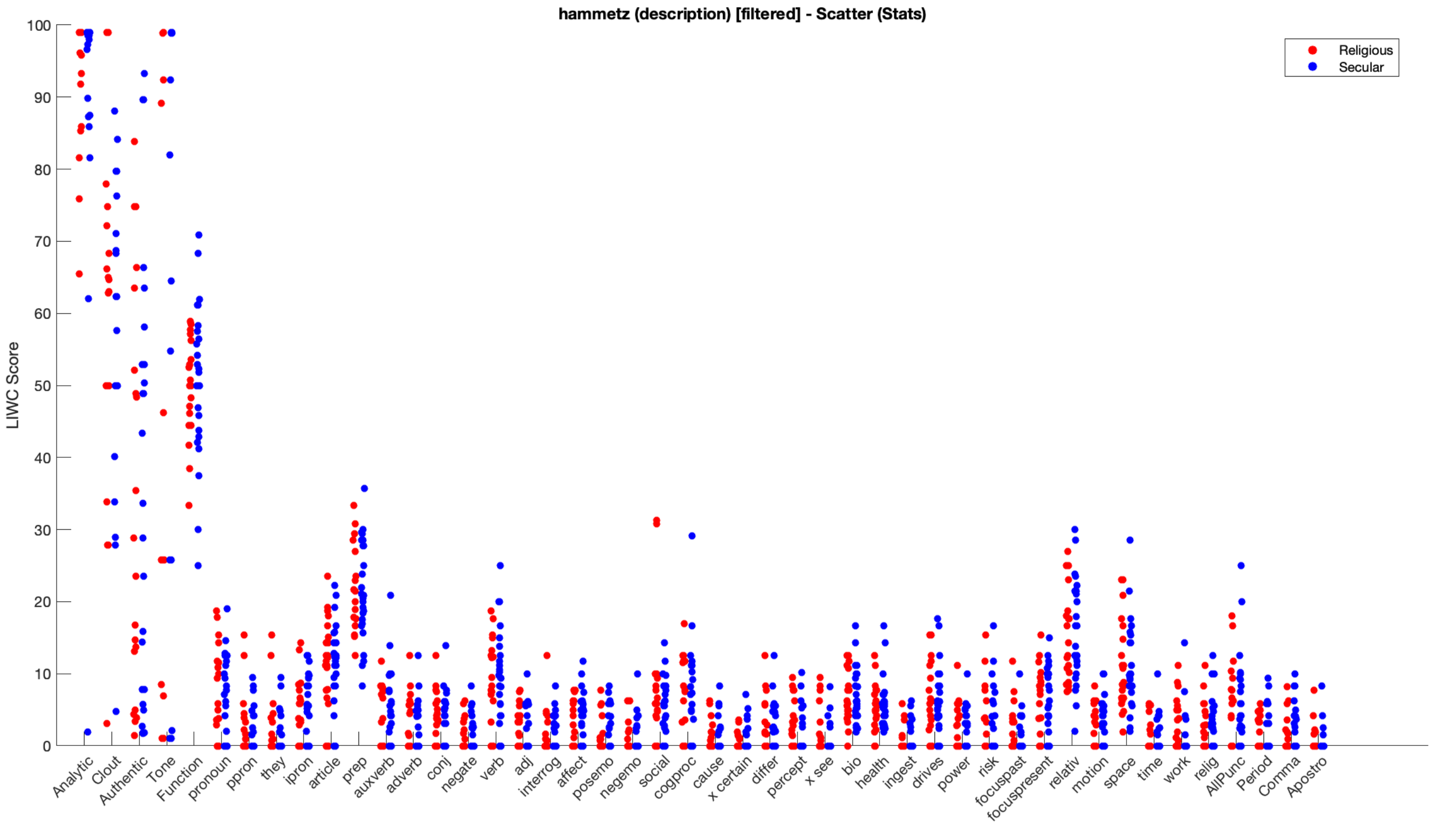


*Fig. S2f – LIWC results for the association question, Passover video*

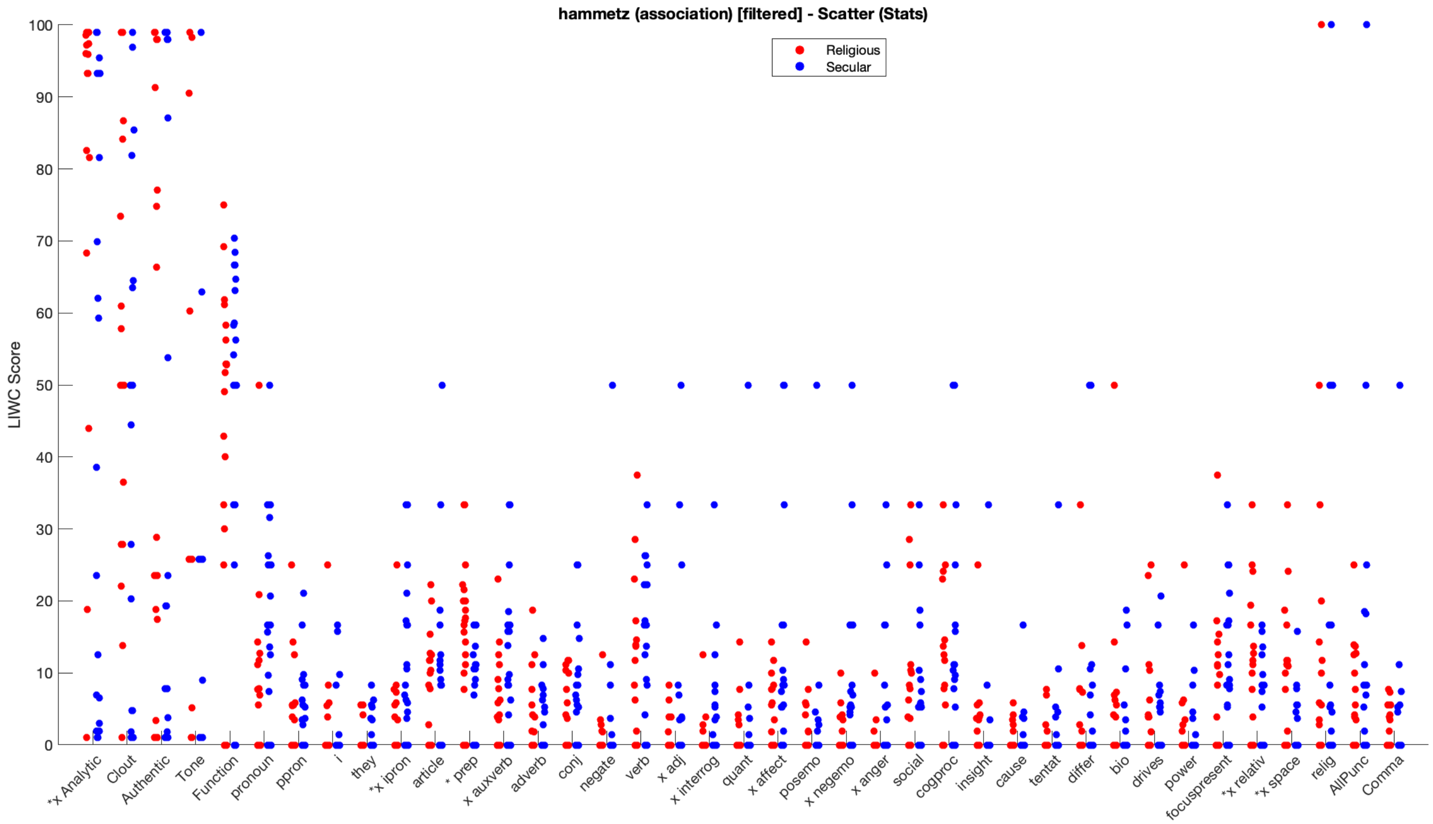

*Cosine Similarity results*

We tested group differences in the cosine similarity of four measures (token, lemma, POS, semantic) in each video and types of question (description and association). In 5 out of 6 the comparisons (two questions, association and description x three video clips, Neutral, Kosher, Passover), no differences were found between the religious and secular in terms of in-group homogeneity in the text responses. Only in the description question ("what did you see in the video?") for the Kosher video-clip, significant differences were found between the groups, with the Religious being more homogenous in the responses in three out of the four measures.

Note: $p < 0.05$ are marked in bold.

*Table S4 – Cosine Similarity statistic results*

| Video | Question type | Measure | *Estimate* | *CI* | *p* |
|---|---|---|---|---|---|
| Neutral | Description | token | -0.01 | –0.04 – 0.02 | 0.559 |
| | | lemma | -0.01 | –0.04 – 0.02 | 0.581 |
| | | pos | 0.12 | –0.12 – 0.36 | 0.329 |
| | | semantic | 0.02 | –0.12 – 0.16 | 0.756 |
| | Association | token | 0.01 | –0.05 – 0.06 | 0.853 |
| | | lemma | 0.00 | –0.05 – 0.06 | 0.955 |
| | | pos | 0.00 | –0.29 – 0.29 | 1.000 |
| | | semantic | –0.11 | –0.27 – 0.05 | 0.164 |
| Kosher | Description | token | -0.02 | –0.05 – 0.00 | 0.102 |
| | | lemma | -0.04 | –0.06 – –0.01 | **0.007** |
| | | pos | -0.28 | –0.56 – –0.01 | **0.041** |
| | | semantic | -0.15 | –0.26 – –0.03 | **0.017** |
| | Association | token | 0.02 | –0.04 – 0.08 | 0.483 |
| | | lemma | 0.02 | –0.03 – 0.07 | 0.472 |
| | | pos | -0.16 | –0.46 – 0.15 | 0.301 |
| | | semantic | -0.10 | –0.24 – 0.04 | 0.172 |
| Passover | Description | token | -0.00 | –0.03 – 0.03 | 0.899 |
| | | lemma | -0.00 | –0.03 – 0.03 | 0.924 |
| | | pos | 0.05 | –0.19 – 0.30 | 0.663 |
| | | semantic | 0.02 | –0.12 – 0.16 | 0.802 |
| | Association | token | 0.01 | –0.05 – 0.08 | 0.664 |
| | | lemma | -0.01 | –0.08 – 0.06 | 0.854 |
| | | pos | -0.22 | –0.53 – 0.08 | 0.152 |
| | | semantic | 0.00 | –0.16 – 0.16 | 1.000 |

### In-group neural synchronization in overlapping regions

*Fig. S3 – In-group neural synchronization in overlapping regions*

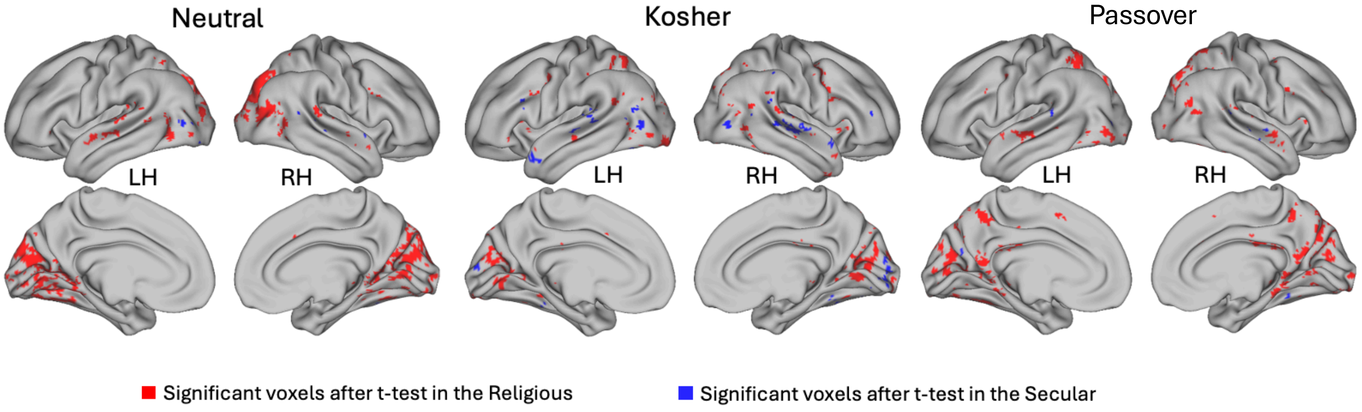


In regions where the in-group Intersubject Correlation (ISC) scores were above chance in both groups, the scores differences were tested using two-tails t-test. Marked in red are voxels in which the Religious' ISC scores were significantly ($p < .05$) higher than the Secular's scores, and in blue – vice versa. In all the videos, there were significantly ($p < .001$) more voxels in which the Religious had higher ISC scores (tested using Chi-squared proportion test) (see table 5).

*Table S5 – comparison of in-group neural synchronization in overlapping regions*

| Video | Overlapping voxels | Significant voxels, Religious > Secular | Significant voxels, Secular > Religious |
|---|---|---|---|
| Neutral | 35686 | 7003 | 283 |
| Kosher | 53291 | 3782 | 1331 |
| Passover | 50090 | 5770 | 406 |

### In-group ISC comparison grouped by Gyri and networks

Note: 0 marks voxels that are not mapped into a gyrus / network in the Brainnetome atlas.

*Table S6a – In-group ISC comparison grouped by gyri*

| Comparison / Network | video | overlap | unique religious | unique secular | unique religious > secular after t-test and FDR | unique secular > religious after t-test and FDR |
|---|---|---|---|---|---|---|
| Amygdala | Passover | 0.008 | 0.059 | 0.007 | 0.015 | 0 |
| | Kosher | 0.102 | 0.269 | 0.049 | 0.063 | 0.01 |
| | Neutral | 0.076 | 0.127 | 0.013 | 0.059 | 0 |
| Basal Ganglia | Passover | 0.002 | 0.104 | 0.013 | 0.069 | 0.002 |
| | Kosher | 0.003 | 0.07 | 0.02 | 0.032 | 0.004 |
| | Neutral | 0.016 | 0.133 | 0.036 | 0.076 | 0.007 |
| Cingulate Gyrus | Passover | 0.145 | 0.377 | 0.013 | 0.237 | 0.001 |
| | Kosher | 0.154 | 0.337 | 0.045 | 0.173 | 0.007 |
| | Neutral | 0.04 | 0.179 | 0.006 | 0.129 | 0 |
| Fusiform Gyrus | Passover | 0.651 | 0.033 | 0.022 | 0.011 | 0.004 |
| | Kosher | 0.637 | 0.029 | 0.029 | 0.008 | 0.008 |
| | Neutral | 0.566 | 0.06 | 0.016 | 0.024 | 0.002 |
| Hippocampus | Passover | 0.041 | 0.095 | 0.041 | 0.057 | 0.009 |
| Hippocampus | Kosher | 0.046 | 0.083 | 0.073 | 0.026 | 0.01 |
| Hippocampus | Neutral | 0.013 | 0.093 | 0.015 | 0.046 | 0.001 |
| Inferior Frontal Gyrus | Passover | 0.139 | 0.233 | 0.052 | 0.099 | 0.005 |
| | Kosher | 0.271 | 0.324 | 0.035 | 0.093 | 0.001 |
| | Neutral | 0.047 | 0.291 | 0.025 | 0.143 | 0.002 |
| Inferior Parietal Lobule | Passover | 0.316 | 0.268 | 0.031 | 0.13 | 0.003 |
| | Kosher | 0.45 | 0.245 | 0.056 | 0.057 | 0.005 |
| | Neutral | 0.189 | 0.097 | 0.102 | 0.034 | 0.015 |
| Inferior Temporal Gyrus | Passover | 0.141 | 0.099 | 0.02 | 0.042 | 0.002 |
| | Kosher | 0.155 | 0.123 | 0.037 | 0.06 | 0.011 |
| | Neutral | 0.091 | 0.063 | 0.023 | 0.02 | 0.002 |
| Insular Gyrus | Passover | 0.059 | 0.181 | 0.021 | 0.085 | 0.002 |
| | Kosher | 0.102 | 0.219 | 0.034 | 0.087 | 0.005 |
| | Neutral | 0.019 | 0.105 | 0.024 | 0.049 | 0.003 |
| MedioVentral Occipital Cortex | Passover | 0.906 | 0.054 | 0.014 | 0.013 | 0 |
| | Kosher | 0.833 | 0.104 | 0.019 | 0.048 | 0.002 |
| | Neutral | 0.879 | 0.092 | 0.007 | 0.06 | 0.001 |
| Middle Frontal Gyrus | Passover | 0.029 | 0.185 | 0.013 | 0.106 | 0.001 |
| | Kosher | 0.131 | 0.14 | 0.048 | 0.03 | 0.007 |
| | Neutral | 0.007 | 0.097 | 0.013 | 0.041 | 0.002 |
| Middle Temporal Gyrus | Passover | 0.233 | 0.152 | 0.028 | 0.076 | 0.004 |
| | Kosher | 0.445 | 0.19 | 0.053 | 0.057 | 0.006 |
| | Neutral | 0.242 | 0.08 | 0.109 | 0.022 | 0.025 |
| Orbital Gyrus | Passover | 0.005 | 0.099 | 0.008 | 0.056 | 0 |
| | Kosher | 0.031 | 0.109 | 0.031 | 0.042 | 0.005 |
| | Neutral | 0.01 | 0.054 | 0.006 | 0.023 | 0.001 |
| Paracentral Lobule | Passover | 0.002 | 0.195 | 0.004 | 0.152 | 0 |
| | Kosher | 0.022 | 0.125 | 0.002 | 0.073 | 0 |
| | Neutral | 0 | 0.017 | 0.002 | 0.007 | 0.001 |
| Parahippocampal Gyrus | Passover | 0.295 | 0.075 | 0.012 | 0.04 | 0.003 |
| | Kosher | 0.137 | 0.07 | 0.037 | 0.019 | 0.006 |
| | Neutral | 0.146 | 0.1 | 0.022 | 0.059 | 0.001 |

| Comparison / Network | video | overlap | unique religious | unique secular | unique religious > secular after t-test and FDR | unique secular > religious after t-test and FDR |
|---|---|---|---|---|---|---|
| Postcentral Gyrus | Passover | 0.061 | 0.191 | 0.041 | 0.113 | 0.003 |
| | Kosher | 0.059 | 0.212 | 0.034 | 0.072 | 0.007 |
| | Neutral | 0.021 | 0.01 | 0.042 | 0.003 | 0.011 |
| Precentral Gyrus | Passover | 0.147 | 0.176 | 0.068 | 0.08 | 0.006 |
| | Kosher | 0.213 | 0.187 | 0.04 | 0.068 | 0.002 |
| | Neutral | 0.056 | 0.161 | 0.038 | 0.054 | 0.005 |
| Precuneus | Passover | 0.414 | 0.451 | 0.008 | 0.322 | 0.002 |
| | Kosher | 0.202 | 0.339 | 0.048 | 0.126 | 0.003 |
| | Neutral | 0.106 | 0.3 | 0.034 | 0.191 | 0.003 |
| Superior Frontal Gyrus | Passover | 0.037 | 0.146 | 0.022 | 0.077 | 0.002 |
| | Kosher | 0.069 | 0.17 | 0.052 | 0.053 | 0.007 |
| | Neutral | 0.011 | 0.123 | 0.024 | 0.053 | 0.005 |
| Superior Parietal Lobule | Passover | 0.462 | 0.319 | 0.031 | 0.215 | 0.001 |
| | Kosher | 0.431 | 0.245 | 0.035 | 0.058 | 0.004 |
| | Neutral | 0.097 | 0.173 | 0.043 | 0.07 | 0 |
| Superior Temporal Gyrus | Passover | 0.538 | 0.084 | 0.03 | 0.032 | 0.004 |
| | Kosher | 0.569 | 0.071 | 0.063 | 0.025 | 0.018 |
| | Neutral | 0.494 | 0.09 | 0.057 | 0.04 | 0.01 |
| Thalamus | Passover | 0.09 | 0.104 | 0.047 | 0.052 | 0.004 |
| | Kosher | 0.041 | 0.117 | 0.024 | 0.042 | 0.005 |
| | Neutral | 0.06 | 0.173 | 0.048 | 0.081 | 0.007 |
| Lateral Occipital Cortex | Passover | 0.929 | 0.039 | 0.004 | 0.014 | 0 |
| | Kosher | 0.909 | 0.054 | 0.004 | 0.031 | 0 |
| | Neutral | 0.852 | 0.095 | 0.011 | 0.057 | 0.001 |
| Posterior Superior Temporal Sulcus | Passover | 0.892 | 0.053 | 0.039 | 0.014 | 0.007 |
| | Kosher | 0.972 | 0.012 | 0.011 | 0.001 | 0.001 |
| | Neutral | 0.781 | 0.075 | 0.089 | 0.017 | 0.013 |

*Table S6b – In-group ISC comparison grouped by 17 networks*

| Comparison / Network | video | overlap | unique religious | unique secular | unique religious > secular after t-test and FDR | unique secular > religious after t-test and FDR |
|---|---|---|---|---|---|---|
| Control A | Passover | 0.171 | 0.238 | 0.033 | 0.114 | 0.001 |
| | Kosher | 0.298 | 0.235 | 0.056 | 0.061 | 0.007 |
| | Neutral | 0.054 | 0.184 | 0.047 | 0.082 | 0.006 |
| Control B | Passover | 0.014 | 0.182 | 0.007 | 0.099 | 0 |
| | Kosher | 0.13 | 0.126 | 0.049 | 0.029 | 0.006 |
| | Neutral | 0.008 | 0.063 | 0.023 | 0.023 | 0.005 |
| Control C | Passover | 0.159 | 0.483 | 0.008 | 0.301 | 0 |
| | Kosher | 0.253 | 0.202 | 0.076 | 0.032 | 0.005 |
| | Neutral | 0.002 | 0.185 | 0.029 | 0.082 | 0.005 |
| Default A | Passover | 0.138 | 0.215 | 0.012 | 0.111 | 0.002 |
| | Kosher | 0.108 | 0.261 | 0.037 | 0.112 | 0.004 |
| | Neutral | 0.044 | 0.087 | 0.042 | 0.046 | 0.008 |
| Default B | Passover | 0.092 | 0.159 | 0.022 | 0.087 | 0.001 |
| | Kosher | 0.208 | 0.218 | 0.051 | 0.065 | 0.007 |
| | Neutral | 0.098 | 0.139 | 0.047 | 0.061 | 0.008 |
| Default C | Passover | 0.562 | 0.186 | 0.019 | 0.109 | 0.003 |
| | Kosher | 0.204 | 0.221 | 0.059 | 0.107 | 0.008 |
| | Neutral | 0.232 | 0.212 | 0.031 | 0.133 | 0.002 |
| Default D (auditory) | Passover | 0.677 | 0.123 | 0.033 | 0.041 | 0.002 |
| | Kosher | 0.793 | 0.141 | 0.02 | 0.042 | 0.002 |
| | Neutral | 0.601 | 0.083 | 0.163 | 0.022 | 0.029 |
| Dorsal Attention A | Passover | 0.815 | 0.083 | 0.019 | 0.043 | 0.001 |
| | Kosher | 0.81 | 0.083 | 0.022 | 0.027 | 0.005 |
| | Neutral | 0.584 | 0.151 | 0.041 | 0.064 | 0.002 |
| Dorsal Attention B | Passover | 0.234 | 0.312 | 0.06 | 0.166 | 0.005 |
| | Kosher | 0.32 | 0.238 | 0.074 | 0.054 | 0.007 |
| | Neutral | 0.074 | 0.164 | 0.054 | 0.064 | 0.003 |
| Limbic-visual Central | Passover | 0.001 | 0.019 | 0.001 | 0.012 | 0 |
| | Kosher | 0.003 | 0.036 | 0.015 | 0.015 | 0.001 |
| | Neutral | 0.001 | 0.009 | 0.001 | 0.003 | 0 |
| Limbic-visual Peripheral | Passover | 0.093 | 0.057 | 0.027 | 0.024 | 0.006 |
| | Kosher | 0.116 | 0.072 | 0.047 | 0.031 | 0.013 |
| | Neutral | 0.05 | 0.053 | 0.03 | 0.024 | 0.007 |
| Salience | Passover | 0.075 | 0.325 | 0.022 | 0.177 | 0.001 |
| | Kosher | 0.241 | 0.287 | 0.05 | 0.091 | 0.007 |
| | Neutral | 0.016 | 0.2 | 0.017 | 0.099 | 0.001 |
| Somatomotor A | Passover | 0.041 | 0.232 | 0.029 | 0.145 | 0.003 |
| | Kosher | 0.037 | 0.238 | 0.017 | 0.094 | 0.001 |
| | Neutral | 0.002 | 0.039 | 0.024 | 0.019 | 0.009 |
| Somatomotor B | Passover | 0.249 | 0.154 | 0.044 | 0.089 | 0.005 |
| | Kosher | 0.246 | 0.125 | 0.05 | 0.043 | 0.014 |
| | Neutral | 0.212 | 0.073 | 0.039 | 0.032 | 0.005 |
| Ventral Attention | Passover | 0.021 | 0.284 | 0.015 | 0.198 | 0.003 |
| | Kosher | 0.082 | 0.279 | 0.019 | 0.131 | 0.002 |
| | Neutral | 0.01 | 0.108 | 0.01 | 0.06 | 0.001 |
| Visual Central | Passover | 0.854 | 0.099 | 0.013 | 0.07 | 0.001 |
| | Kosher | 0.687 | 0.157 | 0.033 | 0.061 | 0.002 |
| | Neutral | 0.677 | 0.226 | 0.005 | 0.158 | 0 |
| Visual Peripheral | Passover | 0.936 | 0.038 | 0.005 | 0.01 | 0 |
| | Kosher | 0.918 | 0.047 | 0.006 | 0.029 | 0.001 |
| | Neutral | 0.91 | 0.051 | 0.012 | 0.024 | 0.001 |

**SVM classification results**

Note: 0 marks voxels that are not mapped into a gyrus / network in the Brainnetome atlas.

*Table S7a – SVM classification results, grouped by gyri*

| Gyrus | prop. In Neutral | prop. In Kosher | prop. In Passover | prop. In religious-videos intersect | prop. In all videos intersect |
|---|---|---|---|---|---|
| Amygdala | 0.018 | 0.000 | 0.000 | 0.000 | 0.000 |
| Basal Ganglia | 0.017 | 0.000 | 0.009 | 0.000 | 0.000 |
| Cingulate Gyrus | 0.076 | 0.089 | 0.154 | 0.114 | 0.043 |
| Fusiform Gyrus | 0.081 | 0.001 | 0.028 | 0.002 | 0.002 |
| Hippocampus | 0.011 | 0.000 | 0.025 | 0.000 | 0.000 |
| Inferior Frontal Gyrus | 0.071 | 0.045 | 0.034 | 0.013 | 0.006 |
| Inferior Parietal Lobule | 0.026 | 0.036 | 0.052 | 0.017 | 0.000 |
| Inferior Temporal Gyrus | 0.019 | 0.017 | 0.008 | 0.000 | 0.000 |
| Insular Gyrus | 0.000 | 0.025 | 0.000 | 0.000 | 0.000 |
| MedioVentral Occipital Cortex | 0.212 | 0.099 | 0.050 | 0.056 | 0.046 |
| Middle Frontal Gyrus | 0.012 | 0.013 | 0.054 | 0.014 | 0.006 |
| Middle Temporal Gyrus | 0.025 | 0.024 | 0.039 | 0.017 | 0.007 |
| Orbital Gyrus | 0.003 | 0.006 | 0.016 | 0.000 | 0.000 |
| Paracentral Lobule | 0.004 | 0.041 | 0.107 | 0.071 | 0.006 |
| Parahippocampal Gyrus | 0.049 | 0.000 | 0.012 | 0.000 | 0.000 |
| Postcentral Gyrus | 0.000 | 0.037 | 0.041 | 0.038 | 0.000 |
| Precentral Gyrus | 0.014 | 0.079 | 0.033 | 0.055 | 0.009 |
| Precuneus | 0.147 | 0.077 | 0.316 | 0.128 | 0.061 |
| Superior Frontal Gyrus | 0.011 | 0.017 | 0.030 | 0.007 | 0.000 |
| Superior Parietal Lobule | 0.017 | 0.063 | 0.169 | 0.052 | 0.021 |
| Superior Temporal Gyrus | 0.018 | 0.028 | 0.021 | 0.011 | 0.000 |
| Thalamus | 0.018 | 0.000 | 0.021 | 0.000 | 0.000 |
| lateral Occipital Cortex | 0.137 | 0.036 | 0.034 | 0.007 | 0.006 |
| posterior Superior Temporal Sulcus | 0.065 | 0.000 | 0.007 | 0.000 | 0.000 |

*Table S7b – SVM classification results, grouped by 17 networks*

| Network | prop. in Neutral | prop. in Kosher | prop. in Passover | prop. in religious-videos intersect | prop. in all videos intersect |
|---|---|---|---|---|---|
| Control A | 0.038 | 0.067 | 0.062 | 0.040 | 0.011 |
| Control B | 0.006 | 0.008 | 0.039 | 0.009 | 0.000 |
| Control C | 0.018 | 0.003 | 0.203 | 0.004 | 0.002 |
| Default A | 0.018 | 0.051 | 0.065 | 0.048 | 0.013 |
| Default B | 0.024 | 0.032 | 0.037 | 0.006 | 0.000 |
| Default C | 0.131 | 0.045 | 0.097 | 0.051 | 0.025 |
| Default D (auditory) | 0.056 | 0.026 | 0.028 | 0.019 | 0.009 |
| Dorsal attention A | 0.092 | 0.026 | 0.048 | 0.020 | 0.010 |
| Dorsal attention B | 0.019 | 0.038 | 0.108 | 0.051 | 0.018 |
| Limbic-visual central | 0.000 | 0.000 | 0.001 | 0.000 | 0.000 |
| Limbic-visual peripheral | 0.005 | 0.008 | 0.001 | 0.000 | 0.000 |
| Salience | 0.035 | 0.034 | 0.077 | 0.032 | 0.011 |
| Somatomotor A | 0.008 | 0.052 | 0.087 | 0.033 | 0.008 |
| Somatomotor B | 0.008 | 0.029 | 0.029 | 0.030 | 0.000 |
| Ventral Attention | 0.018 | 0.042 | 0.089 | 0.073 | 0.032 |
| Visual Central | 0.282 | 0.101 | 0.135 | 0.060 | 0.049 |
| Visual Peripheral | 0.117 | 0.036 | 0.023 | 0.011 | 0.009 |
| 0 | 0.017 | 0.002 | 0.020 | 0.004 | 0.001 |
